# Colorectal adenocarcinomas downregulate the mitochondrial Na^+^/Ca^2+^ exchanger NCLX to drive metastatic spread

**DOI:** 10.1101/2020.05.07.083071

**Authors:** Trayambak Pathak, Maxime Gueguinou, Vonn Walter, Céline Delierneux, Martin T. Johnson, Xuexin Zhang, Ping Xin, Ryan E. Yoast, Scott M. Emrich, Gregory S. Yochum, Israel Sekler, Walter A. Koltun, Donald L. Gill, Nadine Hempel, Mohamed Trebak

## Abstract

Despite the established role of mitochondria in tumorigenesis, the molecular mechanisms by which mitochondrial Ca^2+^ (mtCa^2+^) signaling regulates tumor growth and metastasis remain unknown. The crucial role of mtCa^2+^ in tumorigenesis is highlighted by the altered expression of proteins mediating mtCa^2+^ uptake and extrusion in cancer cells. Here, we demonstrate that expression of the mitochondrial Na^+^/Ca^2+^ exchanger NCLX (*SLC8B1*) is decreased in colorectal tumors and is associated with advanced-stage disease in patients. We reveal that downregulation of NCLX leads to mtCa^2+^ overload, mitochondrial depolarization, mitophagy, and reduced tumor size. Concomitantly, NCLX downregulation drives metastatic spread, chemoresistance, the expression of epithelial-to-mesenchymal transition (EMT), hypoxia, and stem cell pathways. Mechanistically, mtCa^2+^ overload leads to an increase in mitochondrial reactive oxygen species (mtROS) which activates HIF1α signaling supporting the metastatic behavior of tumor cells lacking NCLX. Our results reveal that loss of NCLX expression is a novel driver of metastatic progression, indicating that control of mtCa^2+^ levels is a novel therapeutic approach in metastatic colorectal cancer.

**Highlights:**

- The expression of NCLX is decreased in colorectal tumors and is associated with advanced-stage disease in patients.
- NCLX plays a dichotomous role in colorectal tumor growth and metastasis.
- NCLX downregulation causes mitophagy and reduced colorectal cancer tumor growth.
- NCLX downregulation induces stemness, chemoresistance and metastasis through mtCa^2+^/ROS/HIF1α signaling axis.

**Graphical Abstract:** 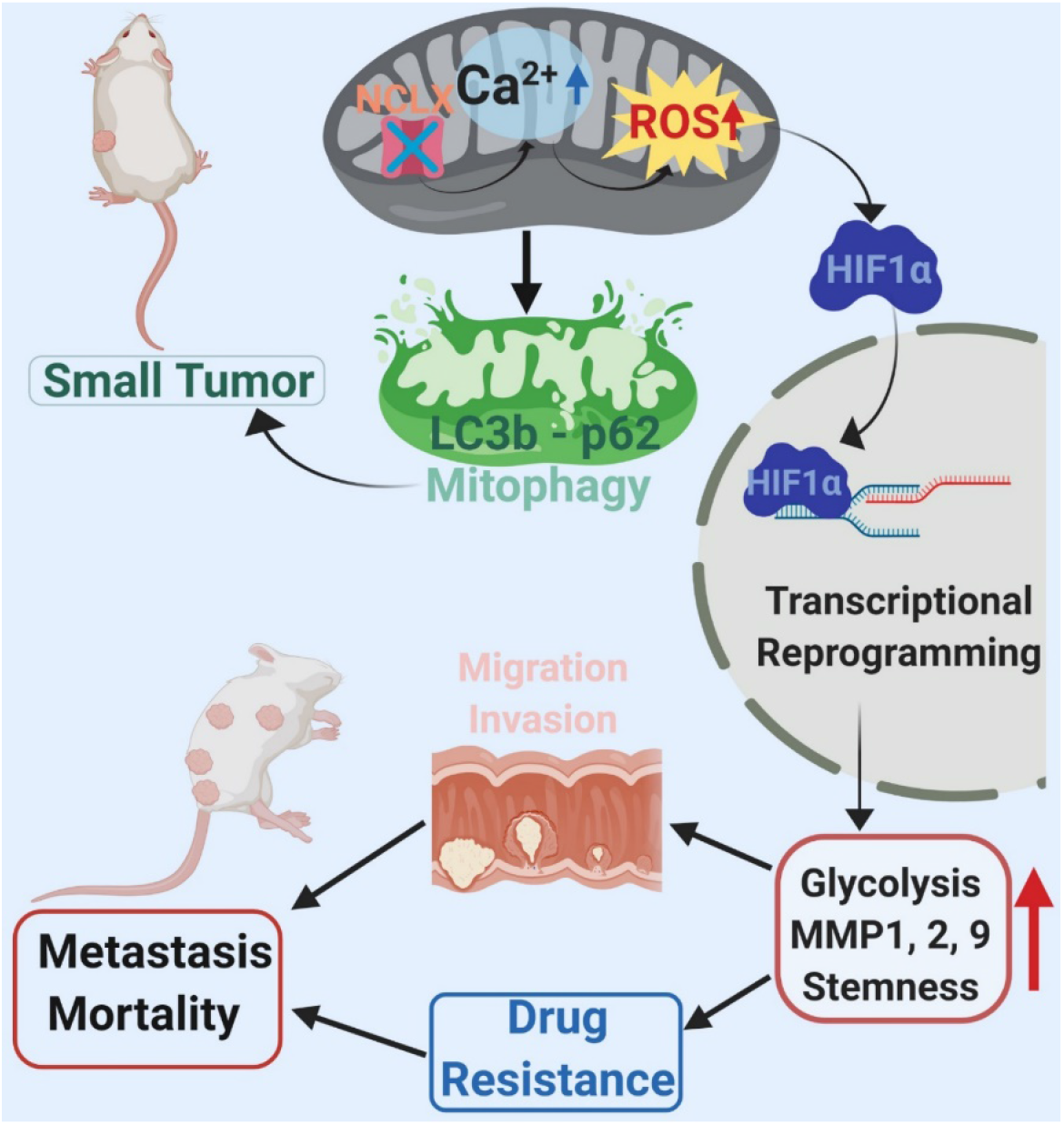

**Significance:** Mitochondrial Ca^2+^ (mtCa^2+^) homeostasis is essential for cellular metabolism and growth and plays a critical role in cancer progression. mtCa^2+^ uptake is mediated by an inner membrane protein complex containing the mitochondrial Ca^2+^ uniporter (MCU). mtCa^2+^ uptake by the MCU is followed by a ∼100-fold slower mtCa^2+^ extrusion mediated by the inner mitochondrial membrane ion transporter, the mitochondrial Na^+^/Ca^2+^ exchanger NCLX. Because NCLX is a slower transporter than the MCU, it is a crucial rate-limiting factor of mtCa^2+^ homeostasis that cannot easily be compensated by another Ca^2+^ transport mechanism. This represents the first study investigating the role of NCLX in tumorigenesis and metastasis. We demonstrate for the first time that colorectal cancers exhibit loss of NCLX expression and that this is associated with advanced-stage disease. Intriguingly, decreased NCLX function has a dichotomous role in colorectal cancer. Thus, we reveal that NCLX loss leads to reduced primary tumor growth and overall tumor burden *in vivo*. Yet, the consequential increases in mtCa^2+^ elicit pro-survival, hypoxic and gene transcription pathways that enhance metastatic progression. This dichotomy is a well-established feature of chemoresistant and recurrent tumor cells including cancer stem cells. Moreover, the downstream changes elicited by NCLX loss are reminiscent of mesenchymal colorectal cancer subtypes that display poor patient survival. Our data indicate that the demonstrated changes to the mtCa^2+^/mtROS/HIF1α signaling axis elicited through the loss of NCLX are a key adaptation and driver of metastatic colorectal cancer.

## Introduction

Mitochondria are adaptable cellular organelles critical for a spectrum of essential functions including ATP generation, cell signaling, cell metabolism, cell proliferation, and cell death (Vyas et al., 2016). This central role causes mitochondria to fulfill crucial role as mediators of tumorigenesis. Mitochondria sense changes in energetic, biosynthesis, and cellular stress, and adapt to the surrounding tumor environment to modulate cancer progression and drug resistance (Tosatto et al., 2016; Vyas et al., 2016). One critical function of mitochondria that is poorly understood in cancer cells is the role of these organelles as a major hub for cellular Ca^2+^ signaling (De Stefani et al., 2016; Pathak and Trebak, 2018). Essentially all mitochondrial functions are controlled by changes in mitochondrial Ca^2+^ (mtCa^2+^) levels. Increased mitochondrial Ca^2+^ uptake stimulates mitochondrial bioenergetics through the activation of Ca^2+^-dependent dehydrogenases of the tricarboxylic acid (TCA) cycle in the mitochondrial matrix (Hansford, 1994; McCormack et al., 1990; Montero et al., 2000). In turn, mtCa^2+^-mediated changes in mitochondrial metabolism can alter the generation of mitochondrial reactive oxygen species (mtROS). In addition, through changes in Ca^2+^ uptake and extrusion, mitochondria shape the spatial and temporal nature of cytosolic Ca^2+^ signals to regulate downstream gene expression programs (De Stefani et al., 2016; Pathak and Trebak, 2018).

Mitochondria form very close contact sites with the endoplasmic reticulum (ER) known as mitochondria-associated membranes (Booth et al., 2016). These contact sites are hotspots of communication through which Ca^2+^ release from the ER via inositol-1,4,5-trisphosphate receptors (IP3R) is efficiently transferred to the mitochondrial matrix through the mitochondrial Ca^2+^ uniporter (MCU) channel complex (Baughman et al., 2011; Booth et al., 2016; De Stefani et al., 2011; Wu et al., 2018). This ensures that the bioenergetic output of the cell is tailored to the strength of cell stimulation by growth factors (Pathak and Trebak, 2018). The major mtCa^2+^ extrusion route is mediated by the mitochondrial Na^+^/Ca^2+^/ Li^+^ exchanger (NCLX) (Palty et al., 2010). The balance between Ca^2+^ uptake by MCU and Ca^2+^ extrusion by NCLX is critical for maintaining mtCa^2+^ homeostasis, which in turn regulates metabolism and cell fate. Perturbations in mtCa^2+^ homeostasis have been linked to a multitude of diseases including cancer (De Stefani et al., 2016; Pathak and Trebak, 2018), and altered mtCa^2+^ homeostasis has recently emerged as a novel hallmark of cancer cells (Danese et al., 2017; Kerkhofs et al., 2018; Paupe and Prudent, 2018). However, the underlying molecular mechanisms by which mtCa^2+^ regulate cancer progression are still poorly understood.

In recent studies, altered MCU expression has been reported in cancer cells, and increased expression of MCU and alterations in proteins that regulate MCU channel activity have been associated with increased mtCa^2+^ and downstream effects that contribute to proliferation and tumor progression (Koval et al., 2019; Marchi et al., 2013; Ren et al., 2017; Tosatto et al., 2016; Vultur et al., 2018). In comparison to MCU, NCLX activity is ~100 fold slower, thus NCLX operation is the rate-limiting factor in mtCa^2+^ homeostasis (Ben-Kasus Nissim et al., 2017; Palty et al., 2010). Cardiomyocyte-specific MCU knockout mice have unaltered levels of mitochondrial matrix Ca^2+^ and no obvious phenotype, whereas cardiomyocyte-specific NCLX knockout mice display mtCa^2+^ overload and die from sudden cardiac arrest (Luongo et al., 2017),. This suggests that although lack of MCU is compensated for by an alternative pathway, the lack of NCLX is not (Pathak and Trebak, 2018). Given the importance of NCLX as the major extrusion mediator of mitochondrial matrix Ca^2+^, it is imperative to understand how aberrant expression of this protein influences mitochondrial and cellular function in cancer. However, to date, the role of NCLX in tumor biology has not been directly investigated.

Colorectal cancer (CRC) is the third most commonly diagnosed cancer type and the third leading cause of cancer deaths in both men and women in the United States (Siegel et al., 2020). Diagnosis at an advanced stage with significant metastatic spread is associated with a 5-year overall survival rate of less than 15%, demonstrating a need to further understand the underlying mechanisms driving CRC metastasis, recurrence, and development of chemoresistance (Siegel et al., 2020). In order to better classify and inform therapeutic interventions the CRC Subtyping Consortium (CRCSC) derived 4 CRC subtypes based on comprehensive molecular signatures including mutation, DNA copy number alteration, DNA methylation, microRNA, and proteomics data to derive four consensus molecular subtypes (CMS). These 4 subtypes differ in their metastatic potential and in survival outcomes of patients (Guinney et al., 2015). For example, CMS4, which is characterized as mesenchymal, is associated with an epithelial-to-mesenchymal transition (EMT) phenotype and poor patient survival statistics. In addition, colorectal cancer cell-intrinsic transcriptional signatures (CRIS), which exclude the contribution of tumor-associated stroma, similarly demonstrate the existence of distinct subtypes based on transcriptional profiling (Dunne et al., 2017; Isella et al., 2017).

Here, we show that the expression of NCLX is significantly downregulated in human colorectal adenocarcinomas. We demonstrate that a loss of NCLX decreases mtCa^2+^ extrusion in CRC cells and that this increase in mtCa^2+^ has important consequences on colorectal tumor cells: NCLX loss 1) inhibits proliferation and primary tumor growth, while 2) enhancing metastasis, and drug resistance, suggesting that a loss of NCLX contributes to CRC metastatic progression. Importantly, decreased NCLX expression leads to transcriptional changes reminiscent of highly metastatic mesenchymal CRC subtypes, including increased expression of genes regulating EMT and cancer stemness, and decreased expression of cell cycle progression mediators. Mechanistically, decreased expression or loss of NCLX results in mtCa^2+^ overload, causing depolarization of mitochondria, mitophagy, and increased mtROS production, which drives ROS-dependent HIF1α protein stabilization and pro-metastatic phenotypes of NCLX-low CRC cells. Thus, we show a novel dichotomous role of NCLX and mtCa^2+^ in cancer, where reduced NCLX function and consequential increases in mtCa^2+^ lead to reduced tumor growth, while driving a mesenchymal phenotype that leads to increased metastasis and drug resistance.

## Results

### NCLX expression is reduced in colorectal tumors

Using the publicly available Cancer Genome Atlas (TCGA) database, we found that NCLX mRNA levels were significantly downregulated in both colon and rectal adenocarcinoma (COADREAD) tumors as compared to the adjacent normal tissue (**Figure 1A**). Consistent with the TCGA data, we observed a substantial reduction to a near loss in NCLX mRNA in colorectal tumor samples isolated from patients undergoing surgery at Penn State University Medical Center as compared to the paired normal adjacent tissues (**Figure 1B**). There was no difference in NCLX mRNA levels between male and female tumor tissues (**Figure S1A**) and between patients bearing mutated and normal proto-oncogene KRAS and phosphoinositide 3-kinases (PI3K) (**Figure 1C**). However, low NCLX expression was associated with TP53 mutations and wild type BRAF tumors (**Figure 1D**). TCGA data analysis from UALCAN (Chandrashekar et al., 2017) revealed that NCLX mRNA was appreciably reduced in CRC patients of all age groups (**Figure S1B**). Both adenocarcinoma and mucinous adenocarcinoma had a significant reduction in NCLX mRNA levels as compared to the normal tissue (**Figure S1C**). Subsequent analysis revealed a significant loss of NCLX in adenomas with malignant transformation from stage I through stage IV (**Figure 1E**). There was a significant reduction in NCLX mRNA level in late-stage (stage III and IV) colorectal tumors as compared to early-stage (stage I and II) tumors from the TCGA database (**Figure 1E, F**), with similar results when we analyzed the patient samples obtained from Penn State University Medical Center (**Figure 1G**). Together, these results show that NCLX expression is significantly downregulated in CRC specimens, and that NCLX loss correlates with late-stage colorectal adenocarcinomas.

**Figure 1:**
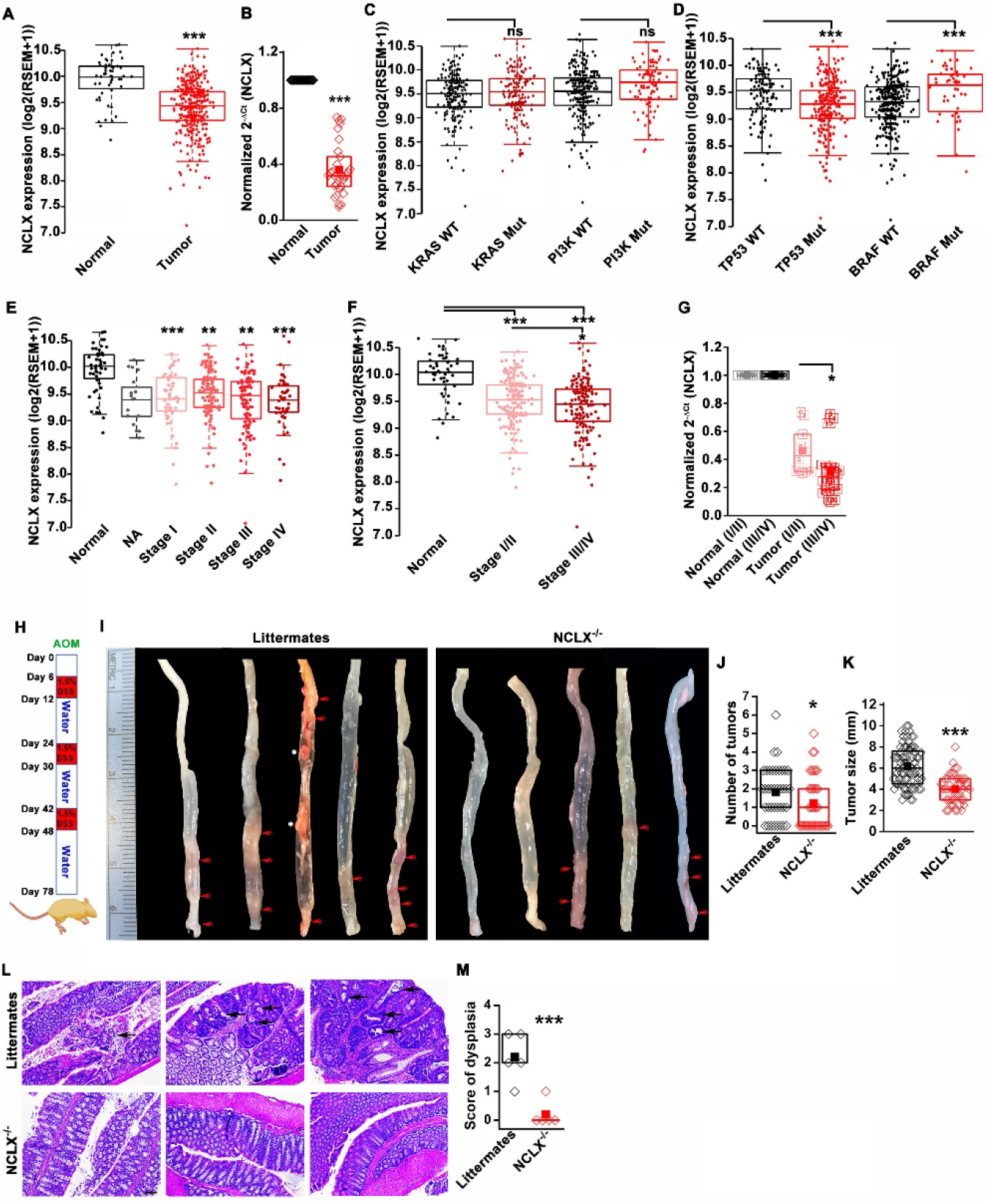
The expression of NCLX, a mtCa^2+^ extrusion mediator in CRC cells, is decreased in CRC tumor samples from human patients. (**A**) TCGA data analysis showing NCLX mRNA levels in tumor tissues and adjacent normal tissues of COADREAD (colon and rectal adenocarcinoma) patients. Each data point represents an individual sample. (**B**) RT-qPCR analysis of NCLX mRNA in tumor tissues (n=30) and adjacent normal tissues (n=30) of CRC patients from Penn State University hospital. (**C**, **D**) TCGA data analysis showing NCLX mRNA level in patients with and without KRAS, PI3K, **(C)** TP53, and BRAF **(D)** mutation. **(E-F)** TCGA data analysis showing NCLX mRNA in tumors at different cancer stages (stages I-IV) **(E)** or combined stage I/II (early stage) and stage III/IV (late-stage) (**F**) of COADREAD tissues compared to adjacent normal tissues. NA= stage not known (**G**) RT-qPCR analysis of NCLX mRNA in combined stage I/II (n=9) and stage III/IV (n=20) CRC tumor samples compared to their adjacent normal tissues obtained from Penn State University hospital. (**H**) Schematic representation of the colitis-associated regimen of AOM and DSS treatment. (**I-K**) Five representative colons from each experimental group are shown **(I),** quantification of the number of tumors (**J**), and tumor volume (**K**) in NCLX^−/−^ and control littermate mice at day 78 after AOM/DSS treatment. The red arrow indicates polyps in the colon and the white star represents fat tissue; n≥30 mice per group. (**L, M**) Three replicates of representative H&E staining of colon sections where black arrows indicate dysplasia (scale bar 500 μm) **(L)**, histology score of dysplasia (scale bar 100 μm) (**M**).Kruskal-Wallis ANOVA was performed to test single variables, between the two groups. *p<0.05, **p<0.01, and ***p<0.001.

### Loss of NCLX has a dichotomous role on tumor growth and metastasis *in vivo*

To determine the functional link between loss of NCLX expression and the development of CRC *in vivo*, we first used global NCLX KO (NCLX^−/−^) mice that were generated using CRISPR/Cas9, where 13 nucleotides (120513241-120513253 on chromosome 13, a region coding for 5 amino acids) were deleted from the first exon of NCLX, resulting in a frameshift mutation and an early stop codon in exon 2 at nucleotide 248 of the coding sequence **(Figure S1D)**. This deletion was confirmed by genotyping of genomic DNA using specific primers for wildtype and knockout alleles (**Figure S1E**). We then used the NCLX^−/−^ mice and their wildtype littermate controls to determine the contribution of NCLX to the development of colorectal tumors in the colitis-associated colorectal cancer model. We subjected NCLX^−/−^ mice (n≥30) and littermate control mice (n≥30) to one intraperitoneal injection of azoxymethane (AOM; 100 μl of 1mg/ml) and three cycles of dextran sodium sulfate (DSS; 1.5%) in drinking water with two weeks of normal water between each DSS cycle (**Figure 1H**). At day 78, mice were sacrificed, and their colorectal tracts were harvested. Colorectal tissues from five representative mice from each experimental group are shown in (**Figure 1I),** and the tissues from fifteen additional mice are depicted in (**Figure S1F, G)**. Although there is a clear association with NCLX loss and CRC in TCGA data, the colons of NCLX^−/−^ mice displayed approximately 50% less tumors than those of littermate control mice (**Figure 1I, J and Figure S1F, G**). Further, the tumors that developed in the colons of NCLX^−/−^ mice were markedly smaller than those in the colons of littermate control mice, as determined by measurements of tumor volume (**Figure 1K**). Histological analyses of colon tissues revealed significantly reduced dysplasia in the colons of NCLX^−/−^ mice compared with colons from littermate control mice (**Figure 1L, M**).

The DSS/AOM administration in mice is an excellent tumor growth model that is not associated with tumor metastases. Therefore, to further investigate the role of NCLX on CRC tumor growth and metastatic spread *in vivo*, we utilized the human CRC cell line HCT116 in an intrasplenic xenograft model, where NCLX was knocked out using the CRISPR/Cas9 system. For *in vivo* studies NCLX KO clone #33 was used, which was generated using a guide RNA (g_1_) which resulted in a single cut at nucleotide 150 in exon 1 causing a frameshift mutation and introduction of a stop codon at position 180 in the NCLX open reading frame (**Figure 2A**). The HCT116 NCLX KO cells and their control HCT116 counterparts were tagged with luciferase and injected (3 × 10^5^ cells/mouse) in the spleens of two groups of NOD-SCID mice for a total of 15 mice per experimental group (**Figure 2A**), and *in vivo* metastasis to the colon and liver was assessed by monitoring luciferase bioluminescence. The mice were injected IP with 100 μl luciferin, and the total flux was measured using the In Vivo Imaging System (IVIS) by exposing the mice for 2 min. There was a significant reduction in the total luciferase bioluminescence flux in mice xenografted with HCT116 NCLX KO cells by comparison to the HCT116 control-injected mice (**Figure 2B, C**), indicating that the loss of NCLX in CRC cells caused reduced tumor growth. The luciferase flux at 2, 4, and 6 weeks from five representative mice per experimental group are shown in (**Figure 2B)** with the remaining 10 mice represented in (**Figure S2)**. Similarly, the primary tumor volumes (at the time of sacrifice, at six weeks) in the spleens of HCT116 NCLX KO-injected mice were significantly reduced compared to control HCT116-injected mice (**Figure 2D, E**), consistent with the *in vivo* tumor growth in the AOM-DSS model. Interestingly, SCID mice injected with the HCT116 NCLX KO cells showed strikingly increased metastasis (**Figure 2B**), specifically to the liver and colon as compared to control HCT116-injected mice at the time of sacrifice (Week 6; **Figure 2F-H**). We did not observe any metastasis to the lung, heart or brain of mice of both groups in this xenograft model. Significantly, the SCID mice with intra-splenic injection of HCT116 NCLX KO cells had reduced overall survival compared to mice injected with control HCT116 cells (**Figure 2I**), suggesting that increased CRC metastasis is the primary cause of lethality in the HCT116 NCLX KO xenograft model. Altogether, our results show that loss of NCLX causes reduced primary tumor growth with increased metastatic progression of colorectal cancer.

**Figure 2:**
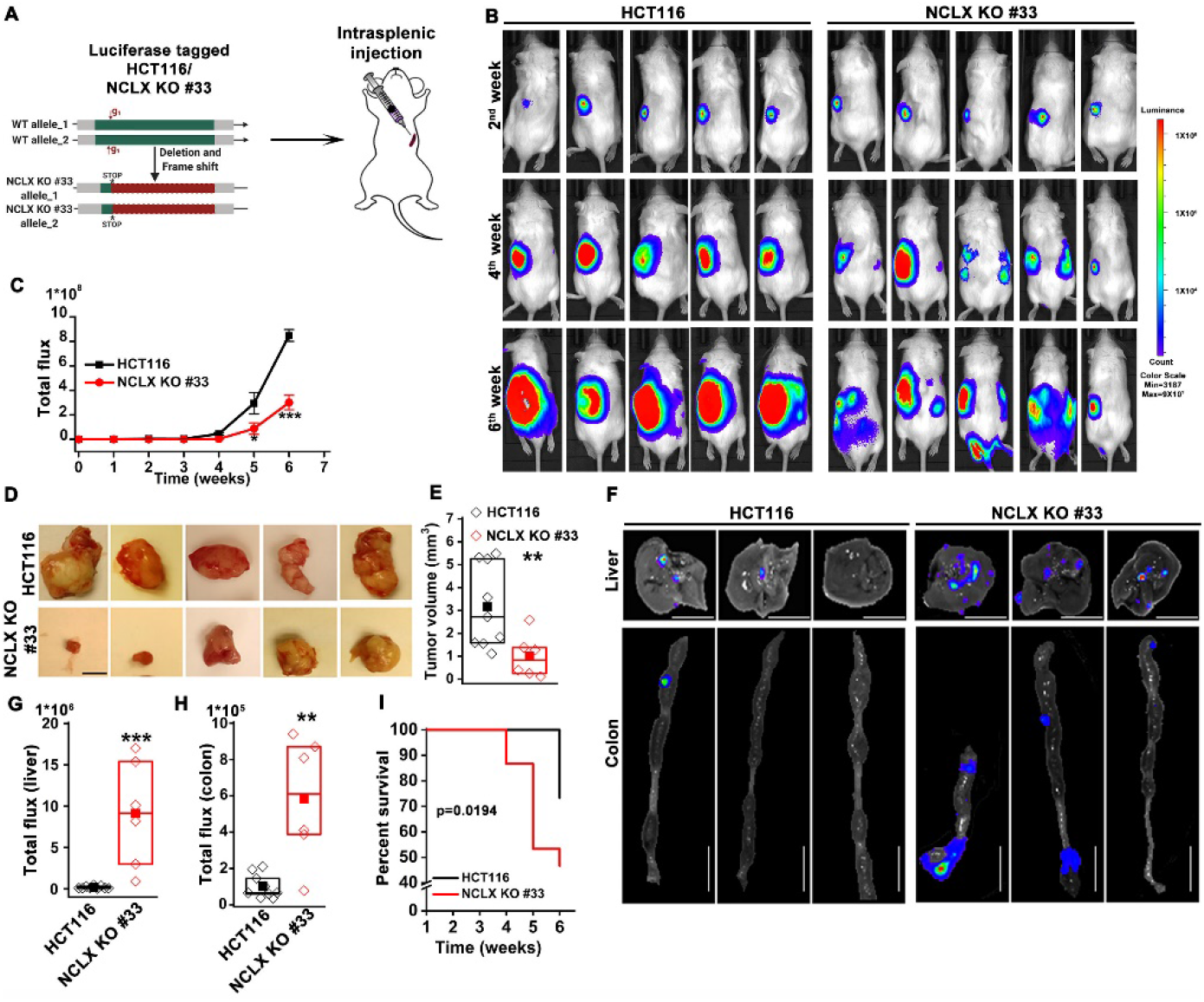
Loss of NCLX has a dichotomous role in tumor growth and metastasis *in vivo*. (**A**) Schematic representation of CRISPR-generated HCT116 NCLX KO #33 cells. Luciferase-tagged control HCT116 and HCT116 NCLX KO #33 cells were injected at 5 × 10^5^ cells/mice into spleens of male NOD SCID mice. (**B-C**) Representative bioluminescence images (5 mice per group) of cancer progression and metastasis in male NOD SCID mice injected with luciferase-tagged HCT116 cells or HCT116 NCLX KO #33 cells **(B)** and quantification of whole-body luciferase count **(C)**. (n =15 mice per group). (**D**, **E**) Representative images of the primary tumors at the site of injection (**D**) and quantification of primary tumor volume at the time of sacrifice (**E**). Scale bar, 5 mm. (**F-H**) Representative image of the liver (scale bar, 5 mm) and corresponding colon (scale bar, 1 cm) (**F**), quantification of luciferase count from the liver (**G**) and the colon (**H**) from NOD SCID mice injected with HCT116 cells or HCT16 NCLX KO #33 cells. (**I**) Survival curve of NOD SCID mice injected with either HCT116 cells or HCT116 NCLX KO #33 cells (*p<0.0194, n =15 mice per group). Kruskal-Wallis ANOVA was performed to test single variables between the two groups.**p<0.01, and ***p<0.001.

### Loss of NCLX inhibits proliferation but enhances migration and invasion of CRC cells

To elucidate the mechanisms by which loss of NCLX elicits these seemingly dichotomous functions on CRC, we investigated the effects of CRISPR/Cas9-mediated knockout, and si/shRNA-driven decreases in NCLX expression in HCT116 and DLD1 CRC cell lines (see Methods and **Figure S3A-H**). To alleviate potential off-target effects of the CRISPR/Cas9 system, we generated several independent clones obtained with three independent guide RNAs (gRNAs; see methods) (**Figure S3A-H**). Genome sequencing and PCR on genomic DNA confirmed NCLX KO **(Figure S3G)**. With the exception of one commercially available polyclonal NCLX antibody (Ben-Kasus Nissim et al., 2017) that is now discontinued, there are currently no reliable NCLX antibodies. All commercially available NCLX antibodies have failed our validation assays and our own attempts to generate a monoclonal antibody against NCLX have not yet produced a reliable clone that detects native levels of NCLX expression. We thus resorted to mRNA quantification, and in all three clones of HCT116 NCLX KO cells, RT-qPCR showed complete absence of NCLX mRNA (**Figure S3D**). Similarly, NCLX KO #06, 24, and 32 of DLD1 cells all had an almost complete deletion of the NCLX open reading frame **(Figure S3E)**, which was confirmed by PCR on genomic DNA (**Figure S3F, G**) and RT-qPCR quantifying mRNA (**Figure S3, H**).

Since the loss of NCLX reduced tumor size in both AOM-DSS and xenograft models **(Figure 1 and 2**), we assessed the effect of reduced NCLX function on the proliferation of CRC cells by CyQUANT proliferation assays. A significant reduction in proliferation was observed in NCLX KO clones of both HCT116 and DLD1 cells (**Figure 3A, B**). To rule out the possibility of long-term compensation in NCLX KO clones, we downregulated NCLX in HCT116 cells using two independent shRNAs and validated the downregulation by qPCR, showing around 60% reduction in NCLX mRNA level in HCT116 cells **(Figure S3I)**. Similarly, transient knockdown of NCLX (NCLX KD) using shRNA reduced HCT116 cell proliferation (**Figure S3J**). The decreases in NCLX KO cell proliferation were not accompanied by appreciable changes in apoptotic cells, although we observed slight increases in cleaved caspase-3 protein using immunofluorescent staining in HCT116 NCLX KO #33 cells as compared to control HCT116 cells, suggesting an increase in apoptosis of NCLX KO CRC cells (**Figure S3K, L**). These data suggest that the reduced tumor sizes observed *in vivo* (**Figures 1 and 2**) due to NCLX knockout are likely a consequence of reduced CRC cell proliferation.

**Figure 3:**
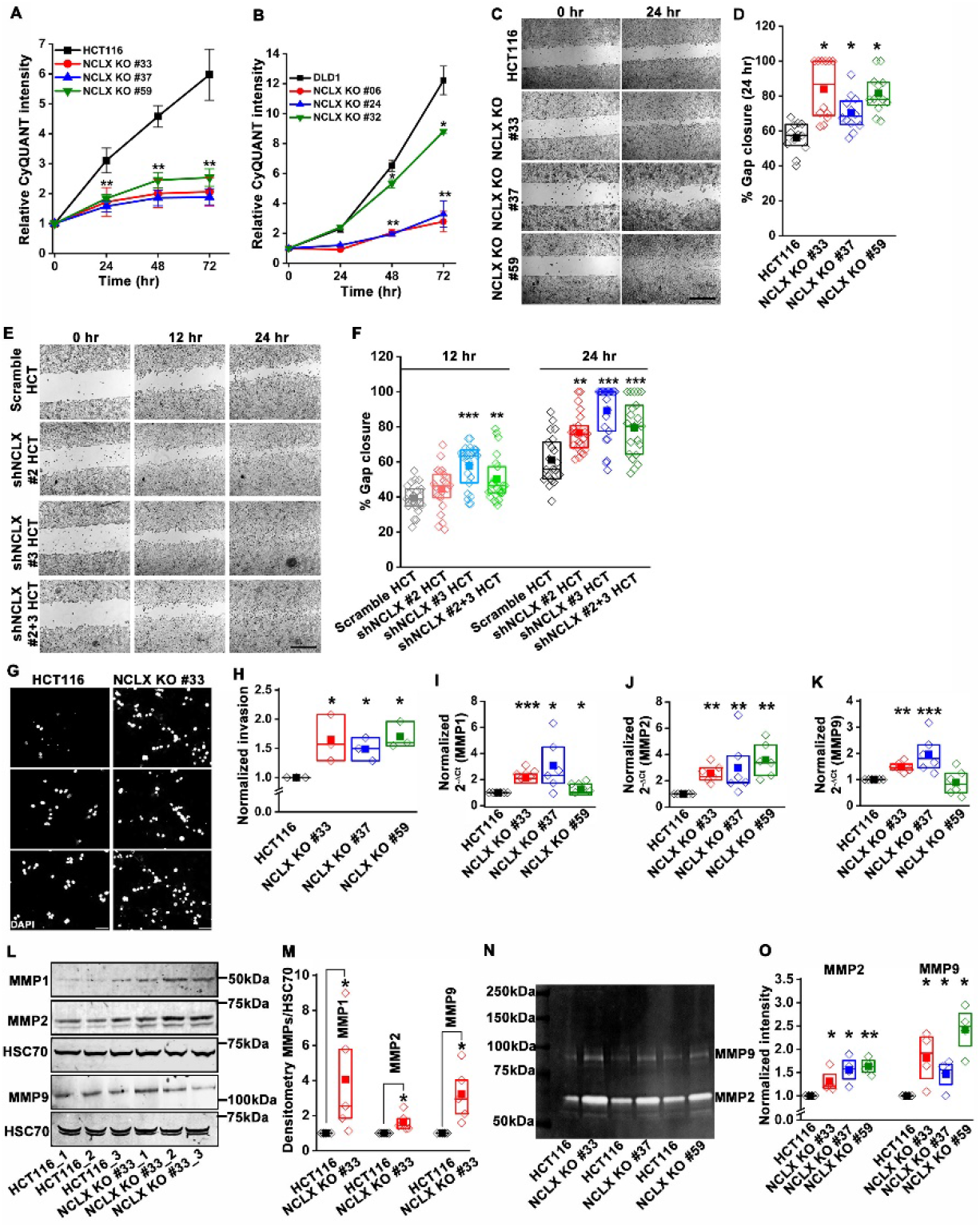
Loss of NCLX inhibits proliferation but enhances migration and invasion of CRC cells. (**A, B**) CyQUANT proliferation assays of HCT116 cells (**A**), DLD1 cells (**B**), and their respective NCLX KO clones. **(C, D)** Representative bright-field images of *in vitro* wound healing migration assay **(C)**, and quantification of % gap closure for HCT116 cells and clones of HCT116 NCLX KO cells after 24 hr **(D)**. Scale bar, 1mm. (**E, F**) *In vitro* migration assays of control HCT116 cells infected with either scramble shRNA or two different shRNA sequences against NCLX **(E)** and quantification of % gap closure (**F**). Scale bar, 1mm. (**G, H**) Representative images of invasion assays performed in triplicates (**G**), and quantification of normalized invasion of HCT116 cells and clones of HCT116 NCLX KO cells **(H)**. Scale bar, 50 μm. (**I-K**) RT-qPCR data showing mRNA expression for MMP1 (**I**), MMP2 (**J**), and MMP9 (**K**) in HCT116 cells and clones of HCT116 NCLX KO cells. (**L**, **M**) Western blot probed with anti-MMP1, anti-MMP2, anti-MMP9, and anti-HSC70 antibody as a loading control (**L**), and quantification of MMPs band intensities normalized to that of HSC70 **(M)**. (**N, O**) Representative gelatin zymogram showing MMP2 and MMP9 activity from HCT116 cells, and their respective clones of HCT116 NCLX KO cells **(N)**, quantification of band intensities of MMP2 and MMP9 activities **(O)**. All experiments were performed ≥three times with similar results. Statistical significance was performed. *p<0.05, **p<0.01 and calculated using one-way ANOVA followed by a post-hoc Tukey test, except for M and O, where the paired t-test was ***p<0.001

Given that NCLX knockout conversely increased metastatic spread in xenograft models (**Figure 2**), we investigated the effect of NCLX knockout and knockdown on the migration pattern of CRC cells. A gap closure assay revealed that all NCLX KO clones of both HCT116 cells (**Figure 3C, D)** and DLD1 cells **(Figure 8N;***light colors*) had a marked increase in migration at 24 hrs. Similar to knockout, shRNA-mediated knockdown of NCLX in HCT116 cells also caused a significant increase in cell migration at 12 and 24 hrs time points (**Figure 3E, F**). Although we showed above that the proliferation of NCLX KO clones of HCT116 cells was inhibited compared to control HCT116 cells **(Figure 3A)**, we ruled out any potential contribution from proliferation in the cell migration assays by analyzing the migration of HCT116 cells and the HCT116 NCLX KO #33 cells at the 6 hr time point. At 6 hr, we observed a significant increase in migration of NCLX KO #33 cells as compared to HCT116 control cells (**Figure S3M**). Effects of proliferation on cell migration were further ruled out by documenting that the increased cell migration of HCT116 NCLX KO cells is preserved in the presence of the cytostatic compound, mitomycin C at 12 hr and 24 hr time points (**Figure S3N**).

Further, the invasive behavior of NCLX KO cells was determined using a Matrigel-coated Boyden chamber assay. A marked increase in invasion was observed in all NCLX KO clones of both HCT116 and DLD1 as compared to their respective controls (**Figure 3G, H, Figure S3O**). Supporting this invasive phenotype are our observations that mRNA levels of the matrix metalloproteinases MMP1, 2, and 9 were significantly upregulated in all the HCT116 NCLX KO clones, as compared to control HCT116 cells (**Figure 3I-K**). Similarly, the protein levels of MMP1, 2, and 9 were also significantly increased in NCLX KO clones of both HCT116 and DLD1 cells (**Figure 3L, M, Figure S3P, Q**). MMP9 and MMP1 protein levels were slightly increased in HCT116 cells in which NCLX was knocked down by shRNA (NCLX KD), although for MMP1, this increase was not statistically significant (**Figure S3R, S**). We then tested the MMPs activity using zymography and revealed that the activity of MMP2 and 9 were markedly increased in all the NCLX KO clones of HCT116 cells compared to control HCT116 cells (**Figure 3N, O**). This was more prominent than changes observed at the mRNA and protein levels, suggesting that the loss of NCLX expression in CRC cells mostly contributes to post-transcriptional regulation of MMPs. Collectively, these results show that the loss of NCLX in CRC cells causes an increase in migration and invasion of CRC cells through increased MMP1, 2, and 9 protein levels and activity. The data also confirm the dichotomous role of NCLX knockout observed *in vivo*, demonstrating that NCLX loss results in decreased cell proliferation and an increase in migratory and invasive phenotypes.

### Loss of NCLX in CRC cells inhibits mtCa^2+^ extrusion, causes mitochondrial perturbations, and enhances mitophagy and mitochondrial ROS

We and others have previously shown that NCLX is the major molecular mediator of mtCa^2+^ extrusion and that inhibiting NCLX expression and function prevents mtCa^2+^ extrusion (Ben-Kasus Nissim et al., 2017; Palty et al., 2010). We measured mtCa^2+^ extrusion in all NCLX KO clones by loading the cells with the mitochondrial Ca^2+^ sensitive dye Rhod-2 AM. Cells were co-loaded with the mitochondrial specific dye MitoTracker green FM as a control. Cells were then stimulated with ATP, a purinergic G protein-coupled receptor (P2Y) agonist that couples to phospholipase Cβ (PLCβ) activation and subsequent inositol-1,4,5-trisphosphate (IP_3_)-dependent release of Ca^2+^ from the ER through IP_3_ receptors (Buvinic et al., 2009; Gonzalez et al., 1989). Upon stimulation with 300 μM ATP in the presence of extracellular Ca^2+^, a portion of Ca^2+^ released from the ER through IP3 receptors is transferred to mitochondria. Hence, cells showed a biphasic response with an increase in Rhod-2 fluorescence followed by a decrease in fluorescence, corresponding to mtCa^2+^ uptake and mtCa^2+^ extrusion, respectively. All NCLX KO clones showed a significant reduction in mtCa^2+^ extrusion (**Figure 4A-H** and **Figure S4A-D**) with no significant change in the rate of mtCa^2+^ uptake. To rule out the possibility of long-term compensation in NCLX KO clones, we transiently downregulated NCLX expression in HCT116 and DLD1 cells using siRNA and validated NCLX knockdown by RT-qPCR. We observed around ~60% reduction in NCLX mRNA levels in both HCT116, DLD1, and HT29 cells (**Figure S4E**). Similar to the NCLX KO cells, HCT116 cells, DLD1 cells, and HT29 cells transfected with siRNA against NCLX (siNCLX) exhibited a significant reduction in mtCa^2+^ extrusion with no significant change in mtCa^2+^ uptake (**Figure S4F-N**). We previously showed that inhibition of mtCa^2+^ extrusion through NCLX knockdown in several cell types leads to the inhibition of plasma membrane ORAI1 channels, reduced Ca^2+^ entry from the extracellular space, and decreased cytosolic Ca^2+^ (Ben-Kasus Nissim et al., 2017). Therefore, we measured cytosolic Ca^2+^ in response to stimulation with ATP in HCT116 cells and their NCLX KO #33 counterparts using the dye Fura-2 and showed a reduction in Ca^2+^ entry in NCLX KO cells with no effect on Ca^2+^ release from the ER (**Figure S4O, P**). Collectively, these results show that inhibition of NCLX function in CRC cell lines leads to enhanced mitochondrial matrix Ca^2+^ concentration due to reduced mtCa^2+^ extrusion and to decreased cytosolic Ca^2+^ due to reduced Ca^2+^ entry across the plasma membrane.

**Figure 4:**
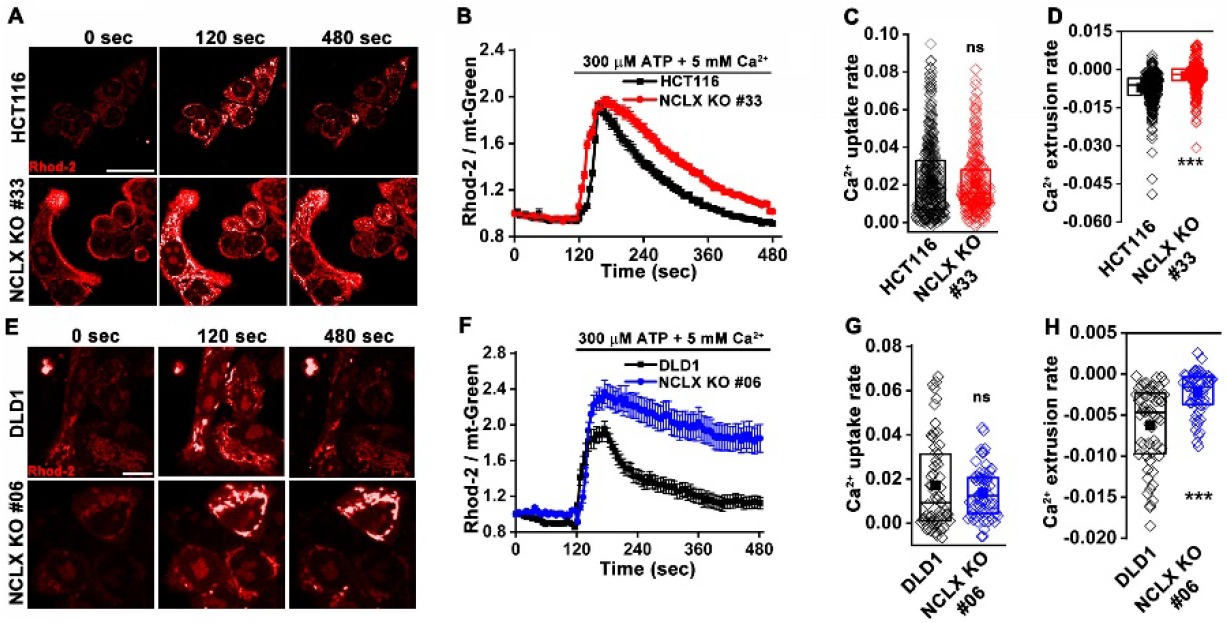
Loss of NCLX in CRC cells inhibits mtCa^2+^ extrusion. (**A**) Representative images of HCT116 cells and HCT116 NCLX KO cells loaded with the mitochondrial Ca^2+^ dye Rhod-2 AM. Cells were stimulated with 300 μM ATP in 5 mM extracellular Ca^2+^ at time 0, and the intensity of Rhod-2 was monitored at different times after ATP stimulation. (**B-D**) The fluorescence ratio of Rhod-2/mt-Green in response to ATP stimulation is monitored as a function of time in HCT116 cells (n=370), and HCT116 NCLX KO cells (n=340) loaded with Rhod-2 and the mitochondrial marker mt-Green (used to normalize the Rhod-2 signal). Stimulation with 300 μM ATP in 5 mM extracellular Ca^2+^ causes rapid mitochondrial Ca^2+^ uptake (ascending phase) followed by Ca^2+^ extrusion (descending phase) (**B**). Quantification of Ca^2+^ uptake rate (**C**) and Ca^2+^ extrusion rate (**D**) from all cells in (**B**). (**E**) Representative images of DLD1 cells and DLD1 NCLX KO cells loaded with the mitochondrial Ca^2+^ dye Rhod-2 AM and stimulated with 300 μM ATP in 5 mM extracellular Ca^2+^ in a manner similar to (**A**). **(F-H)** Similar recordings and analysis as **(B-D)**, except that DLD1 cells (n=63) and DLD1 NCLX KO cells (n=60) were utilized. All the experiments were performed ≥three times with similar results. Statistical significance was calculated using Kruskal-Wallis ANOVA. p*<0.05, **p<0.01, and ***p<0.001

The decrease in mtCa^2+^ extrusion in HCT116 and DLD1 cell clones in which NCLX expression was either reduced by siRNA knockdown or ablated by CRISPR/Cas9 knockout suggested that these clones may be experiencing mitochondrial Ca^2+^ overload. In normal cells, mitochondrial Ca^2+^ overload alters bioenergetics and causes drastic cellular dysfunction leading to cell death (Celsi et al., 2009; Santulli et al., 2015). This occurs mainly through the opening of the mitochondrial permeability transition pore (mPTP) (Bernardi and Di Lisa, 2015; Halestrap, 2009) and subsequent mitochondrial membrane depolarization. **M**) Therefore, the effects of NCLX knockout on mitochondrial membrane potential were measured using the tetramethylrhodamine methyl ester (TMRE) dye. We observed a significant decrease in the accumulation of TMRE in NCLX KO cells, indicating that mitochondria of NCLX KO cells are more depolarized than control HCT116 and DLD1 cells (**Figure 5A, B**).

**Figure 5:**
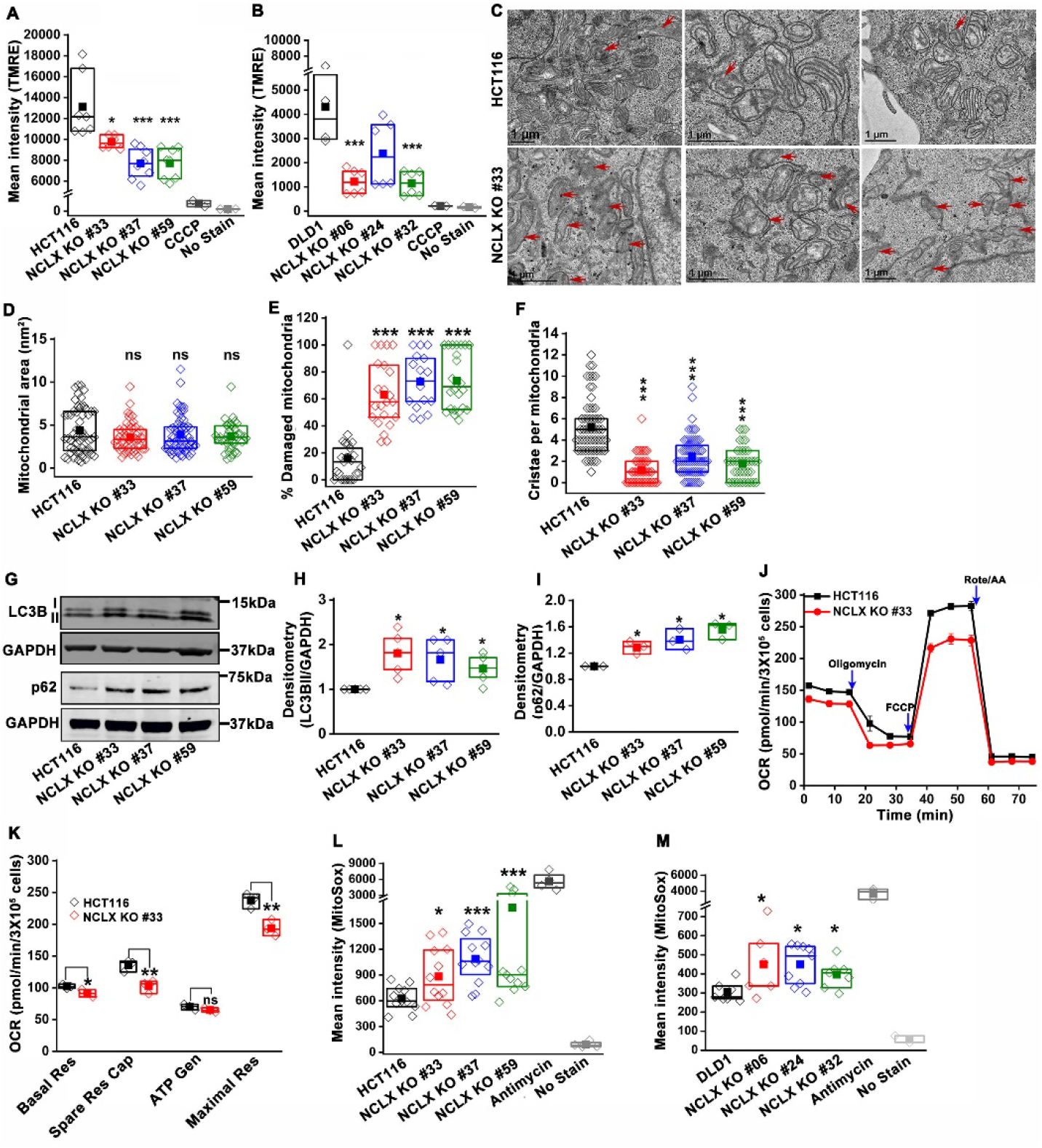
Abrogation of NCLX function in CRC cells causes mitochondrial damage, mitophagy and enhances mitochondrial ROS. (**A**, **B**) Flow cytometric analysis of mitochondrial membrane potential using the dye TMRE in HCT116 cells **(A)** and DLD1 cells **(B)** and their respective NCLX KO clones. Each data point represents one experimental replicate. 100 μM CCCP was added to cells as a positive control, and unstained cells were used as a negative control. (**C**) Representative images of a transmission electron micrograph of HCT116 cells and HCT116 NCLX KO cells. Red arrows indicate damaged mitochondria. (**D-F**) Quantification of mitochondrial area **(D)**, number of damaged mitochondria **(E)**, and number of cristae per mitochondria **(F)** in HCT116 cells and clones of HCT116 NCLX KO cells. (**G-I**) Western blots labeled with indicated antibodies with anti-GAPDH used as a loading control **(G)** and quantification of band intensity of LC3B II (**H**) and p62 (**I**) normalized to GAPDH in HCT116 cells and clones of HCT116 NCLX KO cells. (**J**, **K**) Oxygen consumption rate (OCR) of HCT116 cells and HCT116 NCLX KO #33 cells (**J**), and quantification of basal respiration (Basal Res), spare respiratory capacity (Spare Res Cap), ATP generation (ATP Gen), and maximum respiration (Maximal Res) (**K**). (**L**, Flow cytometric analysis of mitochondrial ROS levels measured with the dye MitoSox in HCT116 cells (**L**) and DLD1 cells (**M**) and their respective NCLX KO clones. As a positive control, 50 μM antimycin was used and unstained cells were used as a negative control. All experiments were performed ≥three times with similar results. Statistical significance was calculated using one-way ANOVA followed by a post-hock Tukey test, except figure H, and I, where paired t-test was used. *p<0.05, **p<0.01, and ***p<0.001

Mitochondrial depolarization is a sign of mitochondrial damage and a major driver of mitophagy, by mediating Pink1 accumulation at the outer mitochondrial membrane (Jin et al., 2010). Examining mitochondrial structure using transmission electron microscopy (TEM) imaging revealed that 60-70% of mitochondria in NCLX KO CRC clones showed altered shape and disrupted cristae compared to control cells (**Figure 5C and Figure S5A-C**). We discovered that mitochondrial membranes and cristae of the NCLX KO clones of HCT116 and DLD1 cells were disrupted (**Figure 5C**, **Figure S5A-C**). Specifically, while overall mitochondrial area was not changed in NCLX KO clones (**Figure 5D**), we observed a greater number of mitochondria with disordered cristae (**Figure 5E**) and a decrease in the number of intact cristae per mitochondrion (**Figure 5F**). We also observed an increase in mitophagic vesicles in NCLX KO cells compared to control HCT116 and DLD1 cells (e.g., see **Figure S5A**). This was accompanied by changes in the autophagy/mitophagy marker LC3BII, which were significantly increased in all NCLX KO clones of HCT116 and DLD1 cells (**Figure 5G, H**, and **Figure S5D, E**). The levels of the p62 protein, a signaling molecule that is downstream of LC3BII (Youle and Narendra, 2010), were also increased in all the NCLX KO clones of HCT116 and DLD1 cells (**Figure 5G, I** and **Figure S5D, E**). Similarly, the shRNA-mediated knockdown of NCLX in HCT116 (see **Figure S3I** for evidence of NCLX mRNA knockdown using shRNA) resulted in increased LC3BII and p62 protein levels (**Figure S5F, G**).

To assess if these mitochondrial perturbations have consequences on mitochondrial electron transport chain function, we measured mitochondrial oxygen consumption rate (OCR) of NCLX KO HCT116 and DLD1 cells using Seahorse extracellular flux assays. While basal and ATP-dependent OCR were slightly decreased, maximal respiration and respiratory reserve capacity were significantly reduced in NCLX KO clones (**Figure 5J, K, and Figure S5H-K**). We determined whether reduced OCR in NCLX KO clones was due to altered protein expression of the mitochondrial respiratory complexes I-V. We did not observe a significant change in the expression of the mitochondrial respiratory complexes I-V between HCT116 cells and their NCLX KO clones **(Figure S5L, M)**. These data suggest that lack of NCLX does not completely abrogate basal mitochondrial respiration, but negatively affects respiratory reserve, which is an indication of the cells ability to enhance mitochondrial respiration in response to higher energy demands.

A further consequence of mtCa^2+^ overload is the generation of mitochondrial reactive oxygen species (mtROS) (Bertero and Maack, 2018; Brookes et al., 2004). In normal cells, enhanced mtROS can lead to mitochondrial dysfunction and cell death; however, many tumor cells utilize mitochondria-derived ROS as cellular signals to drive pro-survival adaptations, including changes in downstream transcription (Vyas et al., 2016). Hence, we measured mtROS in HCT116 and DLD1 CRC cells and their respective NCLX KO clones using the dye MitoSOX and flow cytometry. MitoSOX dye intensity was significantly increased in all the NCLX KO clones of both HCT116 and DLD1 cells (**Figure 5L, M**), indicating increased mtROS in these cells. Using fluorescence microscopy, we also show that downregulation of NCLX with siRNA in HCT116 and DLD1 cells (and in another CRC cell line, HT29; See **Figure S4E** for evidence of NCLX mRNA knockdown in HT29) results in a significant increase in mtROS levels (**Figure S5N, O**). The above data demonstrate that loss of NCLX decreases mtCa^2+^ extrusion, which affects mitochondrial cristae morphology, depolarization of the mitochondrial membrane, induction of mitophagy, decreased respiratory reserve capacity, and enhanced mtROS production.

### Loss of NCLX leads to pro-metastatic transcriptional reprogramming

To further delineate the role ofCLNX loss in CRC, we performed transcriptional profiling of HCT116 cells and their NCLX KO counterparts using RNA sequencing. Gene Set Enrichment Analysis GS(EA) revealed that NCLX knockout drives hte positive enrichment of pathways involved in hypoxia, epithelial to mesenchymal transition (EMT), TGF-β, pron-filammatory, glycolysis, apoptosis and angiogenesis pathways, while negatively influencing gene expression of Myc targets, cell cycle regulation, and oxidative phosphorylation (**Figure 6A-G, and Figure S6A-F**). Thus, GSEA analysis revealed that NCLX loss drives gene expression signatures associated with metastatic progression and inhibition of proliferation, mirroring the phenotypic changes we observed following NCLX knockout and knockdown. Interestingly, these gene expression signatures are shared by the mesenchymal CMS4 CRC subtype, which is characterized by high rates of recurrence, and predictive of poor patient outcome (Guinney et al., 2015).

**Figure 6:**
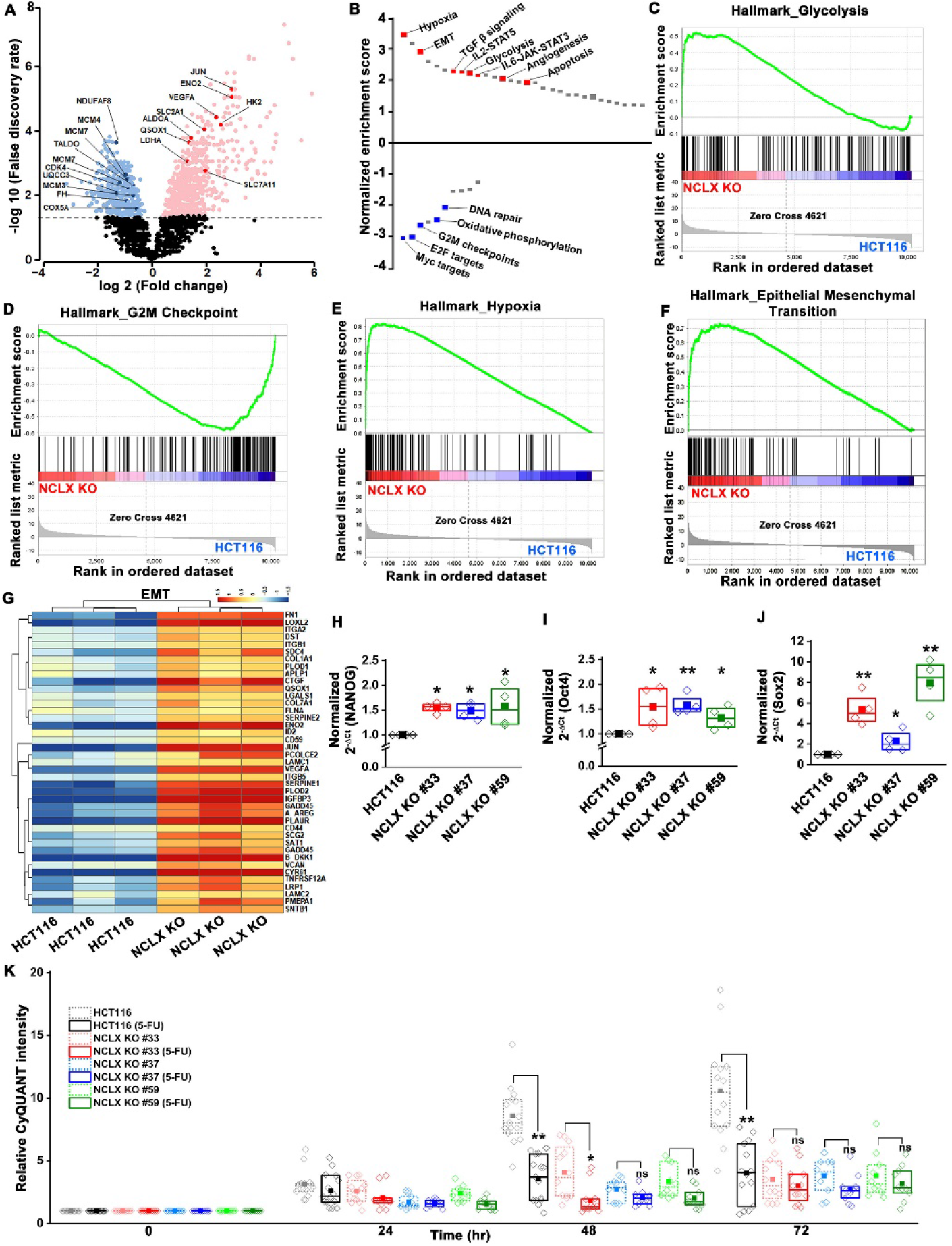
Loss of NCLX leads to pro-metastatic transcriptional reprogramming. (**A**) Volcano plot of differentially expressed genes between HCT116 cells and HCT116 NCLX KO #33 cells showing Log2 fold change vs. false discovery rate. The thresholds in the figure correspond to a false discovery rate < 0.05 and log2 fold change < −1.5 or > 1.5. Genes that are significantly upregulated and downregulated are represented in pink and light blue, respectively. (**B**) Pathway analysis showing normalized enrichment score. Positive enrichment score shows upregulated pathways, and negative enrichment score depicts downregulated pathways. (**C-F**) GSEA analysis of HCT116 cells and HCT116 NCLX KO #33 cells show a positive correlation in the enrichment of hallmark of glycolysis genes **(C)**, a negative correlation of hallmark of G2M checkpoint genes **(D)**, a positive correlation of hallmark of hypoxia-related genes **(E)**, and hallmarks of epithelial to mesenchymal transition genes **(F)**. (**G**) The heat map shows significantly reduced expression of EMT genes in HCT116 NCLX KO #33 cells as compared to control HCT116 cells. (**H-J**) RT-qPCR showing mRNA levels NANOG (**H**), octamer-binding transcription factor 4 (Oct4) (**I**), and SRY-Box Transcription Factor 2 (Sox2) (**J**) in HCT116 cells and clones of HCT116 NCLX KO cells. (**K**) The proliferation of HCT116 cells and HCT116 NCLX KO #33 cells with and without 10 μM 5-FU treatment. All experiments were performed ≥three times with similar results. Statistical significance was calculated using a paired t-test except for K, where one-way ANOVA followed by a post-hock Tukey test was used. *p<0.05, **p<0.01, and ***p<0.001

### NCLX deficiency causes stem cell-like phenotype and chemoresistance of CRC cells

A phenotype of mesenchymal CRC is the enrichment of cancer stem cell traits (Guinney et al., 2015; Polyak and Weinberg, 2009). In agreement with these findings, we found that transcript levels of stem cell markers NANOG, Oct4, Sox2, and Fox3, as well as regulators of the glutathione synthesis pathway implicated in regulating these transcription factors in breast cancer, SLC7A11 and GCLM (Lu et al., 2015), were significantly upregulated in NCLX KO cells (**Figure 6H-J**, **Figure S7A-I**). One exception was the mRNA levels of Oct4 in DLD1 NCLX KO clones, which were not significantly different from those of control DLD1 cells (**Figure S7H**). These data suggest that NCLX KO clones acquire stem cell-like properties.

In most cancer types, a stem cell-like phenotype is associated with enhanced invasion and chemoresistance (Blank et al., 2018; Munro et al., 2018; Reya et al., 2001; Touil et al., 2014). The chemoresistance properties of the NCLX KO clones and respective control HCT116 and DLD1 cells were tested in response to treatment with the antimetabolite agent 5-Fluorouracil (5-FU), which is widely used in the treatment of CRC. First, we performed titration experiments with doses of 5-FU ranging from 1-25 μM. The control HCT116 cells and the HCT116 NCLX KO cells showed a dose-dependent reduction in proliferation in response to 5-FU treatment (**Figure S6G**). The IC_50_ of 5-FU for HCT116 NCLX KO cells (15±1.2 μM) was significantly higher than the control HCT116 cells (8.2±1.2 μM) (**Figure S7J**). Therefore, subsequent experiments were performed with 10 μM 5-FU. The treatment with 10 μM 5-FU caused a significant reduction in proliferation of control HCT116 and DLD1 cells at 24 and 72 hrs (**Figure 6K**, **Figure S7K**). Treatment of the NCLX KO clones of HCT116 (**Figure 6K**) and DLD1 (**Figure S7K**) cells with 10 μM 5-FU yielded only a marginal reduction in proliferation at 72 hrs as compared to non-treated NCLX KO clones, although this reduction was statistically significant for two NCLX KO clones of DLD1 cells at 72 hrs time point (Clone#24 and #32; **Figure S7K**). 5-FU also caused a significant inhibition in migration of control HCT116 cells (**Figure S7L, M**), but did not affect the migratory capabilities of the NCLX KO clones of both HCT116 and DLD1 cells (**Figure S7L, M**).

### NCLX deficiency stabilizes HIF1α and regulates migration in a mtROS-dependent manner

The mesenchymal, invasive, and chemoresistance phenotypes of NCLX KO cells, as well the majority of enriched pathways identified by GSEA, including hypoxia, EMT, glycolysis, angiogenesis, and the suppression of OXPHOS, share a common regulator, the hypoxia-inducible factor HIF1α **(Figure 6A-G, and S6A-F)**. This was intriguing in light of our observations that NCLX knockdown decreases respiration and increases mitochondrial ROS production in CRC (**Figure 5J-M**), as mtROS is a known activator of HIF1α (Bell et al., 2007; Dan Dunn et al., 2015; Hamanaka and Chandel, 2010; Pan et al., 2007). Hence, we measured HIF1α protein levels and found that HIF1α protein levels were strikingly increased in the NCLX KO clones of both HCT116 and DLD1 cells in the absence of a hypoxic stimulus (**Figure 7A-D**). Similarly, HIF1α protein levels were also increased in HCT116 cells in which NCLX was knocked down with shRNA (NCLX KD; **Figure 7E, F**). Importantly, we were able to demonstrate that HIF1α protein stabilization was significantly abrogated in NCLX KO cells when mtROS were scavenged using the mitochondria-targeted antioxidant mitoTEMPO (**Figure 7G, H**). Both AMPK and mTORC1 are known regulators of HIF1α (Dodd et al., 2015; Hudson et al., 2002; Tandon et al., 2011). Therefore, we assessed the phosphorylation levels of AMPK and the ribosomal protein S6K (a readout of mTORC1 activation) in HCT116 and clones of HCT116 NCLX KO cells. Assessment of AMPK and S6K1 phosphorylation revealed no difference between control HCT116 and clones of HCT116 NCLX KO cells (**Figure S7N-Q**), suggesting that mTORC1 does not contribute to the regulation of HIF1α in response to NCLX ablation. Taken together, these results suggest that the major regulator of HIF1α protein levels in NCLX KO CRC cells is mtROS. Further, a significant reduction in migration was observed when HCT116 NCLX KO cells were treated with mitoTEMPO (**Figure 7I, J**), while this had no effect on the proliferation of either control HCT116 cells or their NCLX KO counterparts (**Figure 7K**).

**Figure 7:**
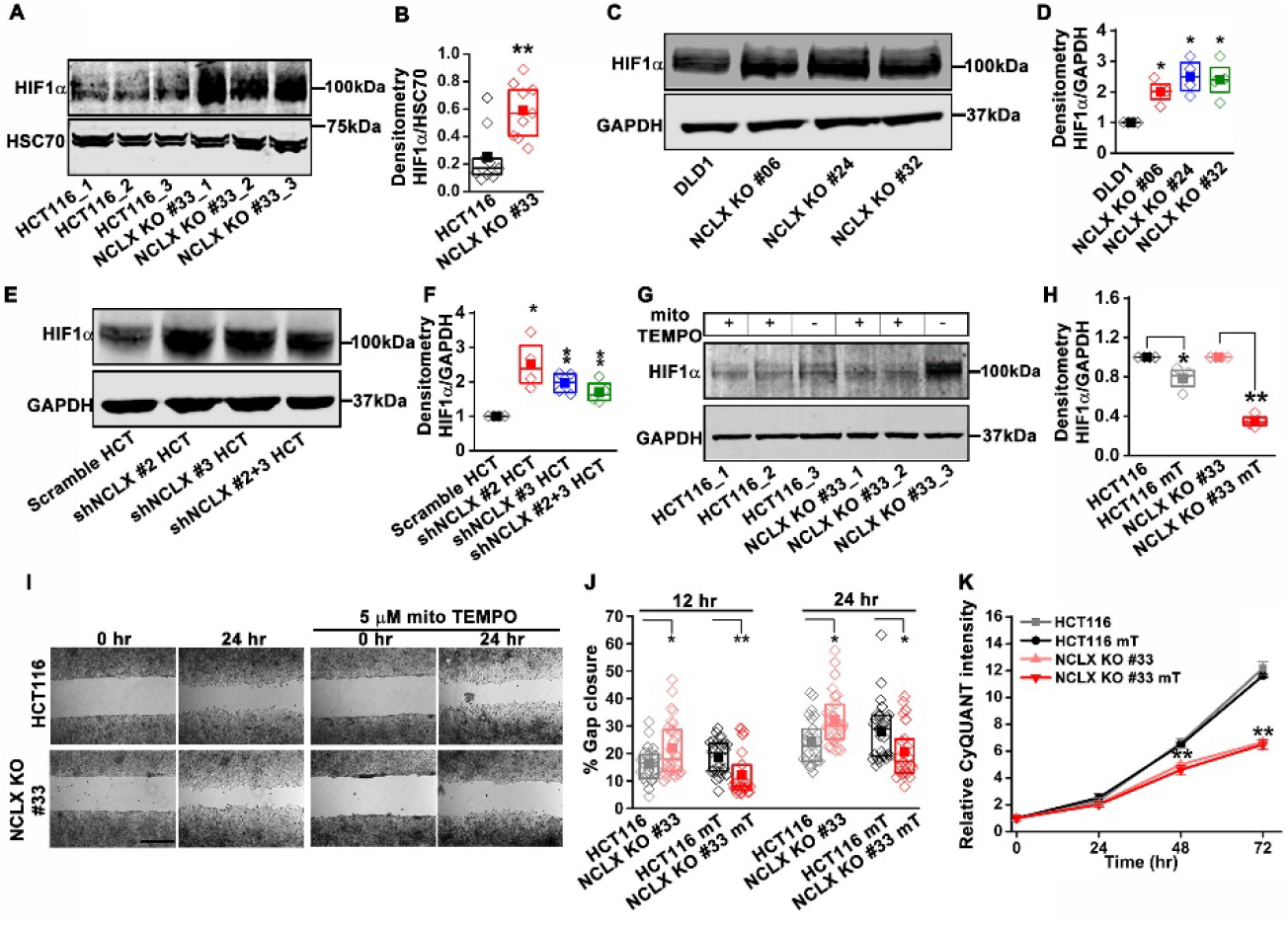
NCLX deficiency stabilizes HIF1α and regulates migration in a mtROS-dependent manner. (**A**-**F**) Western blot of HCT116 cells and clones of HCT116 NCLX KO cells **(A),** DLD1 cells and clones of DLD1 NCLX KO cells **(C)** and HCT116 cells infected with either scramble shRNA or three combinations of shRNA against NCLX **(E)** probed with anti-HIF1α, and anti-HSC70 antibody as a loading control, and quantification of HIF1α band intensity relative to HSC70 (**B, D, and F**). (**G, H**) Western blot showing HIF1α proteins in HCT116 cells and clones of HCT116 NCLX KO cells after treatment with 5 μM mitoTEMPO overnight **(G)**, and quantification of HIF1α band intensity normalized to GAPDH **(H)**. (**I, J**) Representative bright-field images of *in vitro* migration assay **(I)**, and quantification of % gap closure **(J)** of HCT116 cells and clones of HCT116 NCLX KO cells, with and without treatment with 5 μM mitoTEMPO. Scale bar, 1mm. (**K**) Proliferation of HCT116 cells and HCT116 NCLX KO #33 cells with and without 5 μM mitoTEMPO. All experiments were performed ≥three times with similar results. Statistical significance was calculated using the paired t-test unless mentioned otherwise. *p<0.05, **p<0.01, and ***p<0.001

### Glycolysis is critical for migration of NCLX-deficient colorectal cancer cells

HIF1α is an important regulator of glycolysis. The transcriptomic analysis (**Figure 6A** and **Figure S6A**), GSEA pathway, and enrichment analysis (**Figure 6B, C**) showed that glycolysis-related genes were upregulated in HCT116 NCLX KO cells. RT-qPCR results confirmed that the glycolysis-related genes GLUT1 (glucose transporter 1 or SLC2A1), HK2 (hexokinase 2), ALDOA (aldolase A), ENO1 (enolase 1), and LDHA (lactate dehydrogenase A) were distinctly upregulated in all the NCLX KO HCT116 clones (**Figure 8A**). Loss of NCLX in either HCT116 or DLD1 increased the protein levels of HK2, ALDOA, and LDHA (**Figure 8B, C**, and **Figure S8A-D**). We also validated this further using shRNA-mediated knockdown of NCLX (NCLX KD) in HCT116 cells. NCLX mRNA levels were reduced by 60%-70% in NCLX KD cells compared to cells transfected with shRNA scramble control (**Figure S3I**). Similar to NCLX KO, the HCT116 NCLX KD cells showed a modest but significant increase of HK2, ALDOA, and LDHA protein levels (**Figure S8E, F**). Interestingly, we observed a significant downregulation of the pentose phosphate pathway genes in NCLX KO HCT116 cells, including decreased mRNA expression of the enzymes glucose 6-phosphate phosphogluconate dehydrogenase (G6PD), 6-dehydrogenase (PGD) and transketolase (TKT) (**Figure S8G**).

Since inhibition of NCLX function caused upregulation of glycolytic genes, we used Seahorse extracellular flux analysis to determine whether NCLX knockout affects the extracellular acidification rate (ECAR) of HCT116 and DLD1 cells. Consistent with the upregulation of glycolytic genes, ECAR, and glucose dependency were increased in all NCLX KO clones (**Figure 8D-F** and **Figure S8H**). Furthermore, direct measurements of glucose and lactate in growth media using the YSI system also revealed that the glucose consumption and lactate production were markedly increased in NCLX KO clones of both HCT116 and DLD1 cells (**Figure 8G-J**). These data suggest that NCLX KO cells compensate for their decreased respiratory reserve capacity (**Figure 5J, K**, and **Figure S5H-K**) by enhancing glycolytic pathways to meet their energy demands.

**Figure 8:**
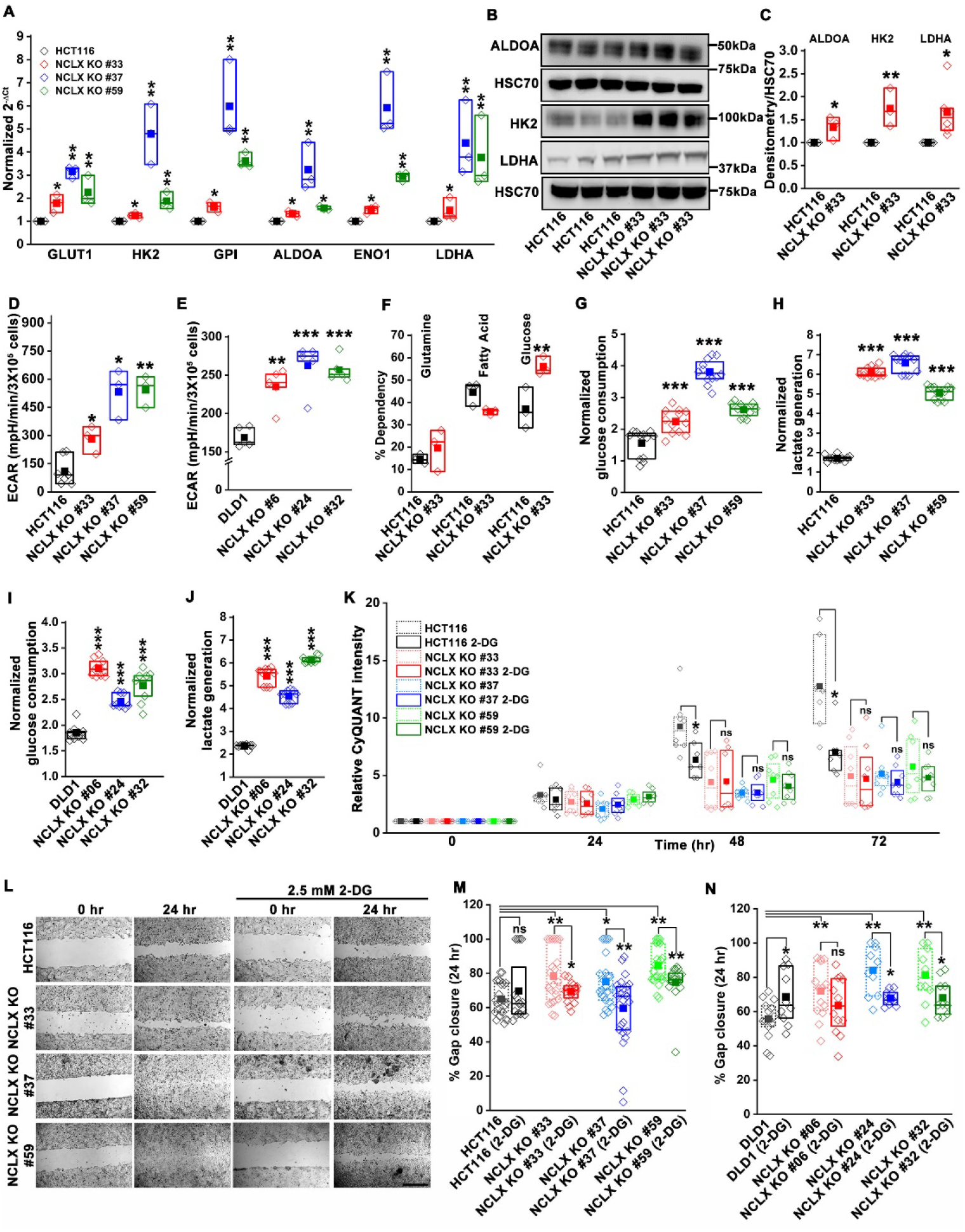
Glycolysis is critical for migration of NCLX-deficient colorectal cancer cells. (**A**) RT-qPCR analysis of glycolytic genes expression in HCT116 cells and three clones of HCT116 NCLX KO cells. RT-qPCR results are plotted as 2^−ΔCt^ relative to tubulin and normalized to control. (**B**, **C**) Representative Western blots probed with anti-ALDOA, anti-HK2, and LDHA antibodies in HCT116 cells and HCT116 NCLX KO #33 cel.lsAnti-HSC70 antibody is used as a loadincgontrol (**B**) and quantification of band intensity relative to that of HSC70 (**C**). (**D, E**) ECAR of HCT116 cells **(D)** and DLD1 cells **(E)** and their respective NCLX KO clones were measured with seahorse using the glycolysis stress test, as described in methods. (**F**) The percentage of metabolite dependency of HCT116 cells and HCT116 NCLX KO #33 cells measured using the Mitochondrial Fuel Flexibility, Dependency, and Capacity test, as described in methods. (**G**-**J**) Measurement of gl ucose consumption and lactate generation in HCT116, clones of H CT116 NCLX KO **(G, H)**, DLD1, and clones of DLD1 NCLX KO **(I, J)** cells normalized to the amount of protein. (**K**) Proliferation of HCT116 cells and clones of HCT116 NCLX KO cells in the presence and absence of 2.5 mM 2-DG. (**L, M**) *In vitro* migration **(L)**, and quantification **(M)** of HCT116 cells and clones of HCT116 NCLX KO cells in the presence and absence of 2.5 mM 2-DG for 24 hr (scale bar, 1mm). All experiments were performed ≥three times with similar results. Statistical significance was calculated using paired t-test, except for K, M, and N, where one-way ANOVA followed by a post-hock Tukey test was used. *p<0.05, **p<0.01, and ***p<0.001

In light of these findings, the glycolytic inhibitor 2-deoxy-D-glucose (2-DG) was used to test if CRC cells that lack NCLX expression are more vulnerable to glycolysis inhibition. While 2-DG caused a significant reduction in proliferation of both DLD1 control and NCLX KO cells (**Figure S8I**), it caused only a mild non-significant reduction in proliferation of NCLX KO clones of HCT116 (**Figure 8K**). However, the enhanced migration of NCLX KO clones was abrogated when glycolysis was inhibited using 2-DG (**Figure 8L-N**), suggesting that increased glycolysis is supporting migration of NCLX KO clones and that targeting this metabolic adaptation may be an avenue to block metastatic progression of CRC cells with low NCLX expression.

## Discussion

In addition to the critical role mitochondria play in cellular bioenergetics, lipid metabolism, and cell death, recent attention has focused on their direct and indirect contributions to mediating crucial signal transduction pathways that control the shift in cellular metabolic activity and changes in gene transcription (Tennant et al., 2009; Tennant et al., 2010). Mitochondria are generators and active participants in Ca^2+^ and ROS signaling (Hempel and Trebak, 2017; Vyas et al., 2016). Growing evidence has shown a critical role of mtCa^2+^ homeostasis in mitochondrial function and cell survival (Celsi et al., 2009; Santulli et al., 2015) and reported a strong correlation between altered mitochondrial function and disease including cancer progression (Porporato et al., 2018). Mitochondria are major mediators of the shift in metabolic activity of cancer cells. This metabolic shift is a mechanism of adaptation to stressors within the tumor microenvironment that is required for cancer cell survival (Vyas et al., 2016). Nevertheless, the mechanisms of mtCa^2+^ in driving mitochondrial signaling and metabolic shifts have largely remained unknown.

Previous studies on the mitochondrial Ca^2+^ uniporter (MCU) indicated that the level of mtCa^2+^ is an important determinant in the progression of different cancer types (Marchi et al., 2013; Tosatto et al., 2016). The downregulation of MCU and subsequent reduction of mtCa^2+^ uptake correlated with resistance to apoptosis of colorectal cancer cell lines (Marchi et al., 2013). Conversely, in triple-negative breast cancer cell lines, the downregulation of MCU caused a reduction in mtROS, HIF1α levels, cell migration, invasion and inhibited metastasis to the lung in xenograft experiments (Tosatto et al., 2016), suggesting that although mtCa^2+^ overload is pro-apoptotic, it is likely beneficial for invasion and metastasis of cancer cell clones that have evaded apoptosis. Similarly, our TCGA data analysis of colorectal cancer patients showed that NCLX mRNA levels are significantly reduced in both early and late-stage tumors with a more pronounced reduction in late-stage tumors, suggesting that mtCa^2+^ overload through reduced NCLX function has a critical role in CRC progression.

It is well established that mtCa^2+^ overload is detrimental to mitochondrial function and is a precursor of death in normal cells through the opening of the mitochondrial permeability transition pore, cytochrome C release from mitochondria and subsequent activation of the caspase family of pro-apoptotic proteases (Giorgi et al., 2012; Pinton et al., 2008). However, the exact mechanisms by which cancer cells adapt and survive under these conditions are not fully understood. Our data show that downregulation or complete loss of NCLX in CRC cells causes mtCa^2+^ overload, an increase in mtROS production and mitochondria depolarization. Our transmission electron microscopy data show that loss of NCLX in CRC cells causes altered mitochondria shape and morphology with disrupted cristae and inner mitochondrial membrane structures. We also show that this coincides with increased LC3B- and p62-dependent mitophagy and decreased cell cycle-related gene expression. This phenotype is consistent with the reduced proliferation of NCLX KO and NCLX KD CRC cells, and the reduced tumor burden and tumor size in NCLX^−/−^ mice subjected to the colitis-associated CRC model. Similarly, our xenograft model yielded smaller primary tumors when HCT116 NCLX KO cells were injected in SCID mice by comparison to xenografts of control HCT116 cells. Thus, the reduction in NCLX expression likely limits proliferation and primary tumor growth. Yet, it is also apparent that clones of cancer cells survive this loss of NCLX and coopt the downregulation of NCLX to enhance pro-survival mitophagy and undergo metabolic reprogramming and gene expression changes that support tumor migration, invasion and metastasis, by initiating a mitochondrial Ca^2+^/ROS signaling axis to drive HIF1α activation.

While it is generally thought that mtCa^2+^ induces mtROS production through increased activation of TCA cycle proteins and consequential increases in ETC function, it is also evident that defective electron transport chain function results in mtROS production (Starkov, 2008). Our data clearly demonstrate that increased mtROS production is accompanied by mitochondrial structural perturbations, decreased OXPHOS, and mitochondrial membrane depolarization. These phenotypes of mitochondrial dysfunction are commonly associated with apoptosis initiation. However, NCLX knock-out resulted in the selection of clones that initiate pro-survival and pro-metastatic adaptations. Mitochondrial depolarization and mtROS production are known initiators of mitophagy (Ashrafi and Schwarz, 2013; Fan et al., 2019; Frank et al., 2012; Schofield and Schafer, 2020; Twig and Shirihai, 2011). Moreover, HIF1α, which is regulated in a mtCa^2+^/mtROS-dependent manner in response to NCLX loss in CRC (**Figure 7K, L**), also contributes to mitophagy, by upregulating the pro-mitophagy proteins BNIP3 and NIX (Bellot et al., 2009; Chourasia et al., 2015; Zhang et al., 2008; Zhang and Ney, 2009).

The gene expression signatures and phenotypes observed in response to NCLX downregulation largely mirror those of the mesenchymal CRC subtype labeled as CMS4. Shared pathways include the increase in EMT, TGF-β, matrix remodeling, stemness, and decreases in Myc and cell cycle progression (Guinney et al., 2015). No single mutation is solely associated with one of the CMS or CRIS subtypes, and at present, the underlying drivers of colorectal cancer molecular subtypes still remain to be fully elucidated (Guinney et al., 2015). However, it is interesting to note that ~ 50% of CMS4 tumors display TP53 mutations and generally lack mutations in BRAF, and that loss of NCLX is statistically linked to TP53 mutant tumors and wild type BRAF CRC tumors (**Figure 1D**). We used two different CRC cell lines HCT116 and DLD1; both of these cell lines are derived from male CRC patients carrying a mutation in KRAS and PIK3CA. TCGA data analysis showed no difference in NCLX expression between CRC patients bearing the wildtype KRAS and PIK3CA and their respective mutations. Interestingly, we also saw lower basal NCLX expression in DLD1 cells that have mutated TP53 (S241F), compared to HCT116 cells, which are wildtype for TP53 (**Figure S8J** compare to **Figure 1D**). Whether increased mtCa^2+^ and subsequent mtROS signaling is one of the underlying drivers of CMS4 tumors remains to be determined. We considered studying the effects of NCLX overexpression on metastasis of HCT116 and DLD1 cells. However, these studies are not feasible because overexpressed NCLX localizes not only to mitochondria but also to the ER and cytosol of these CRC cells, making the interpretation of results of these experiments tenuous.

A metabolic shift towards aerobic glycolysis is a hallmark of cancer progression (Burns and Manda, 2017; Liberti and Locasale, 2016). Our data demonstrated that the loss of NCLX leads to reduced oxygen consumption, increased glycolysis, and increased transcription of major glycolytic enzymes **(Figure 5J, 6B, C and 8)**. This increased transcription of glycolytic enzymes is commonly controlled by HIF1α, which is upregulated in response to increased mtROS resulting from loss of NCLX and subsequent increase in mtCa^2+^. Furthermore, we show that inhibiting glycolysis of NCLX KO CRC cells partially normalizes the increased migration of these cells, consistent with previous findings that showed a critical role of glycolysis in cancer metastasis (Bu et al., 2018; Gillies et al., 2008; Han et al., 2013).

Although the increase in HIF1α protein levels was shown to be critical for the migration and metastasis of cancer cells (Lehmann et al., 2017; Masoud and Li, 2015; Semenza, 2003; Wigerup et al., 2016), the mechanisms by which mtCa^2+^ regulates HIF1α protein levels are not clear. Here, we provide evidence that reduced NCLX function, causing reduced mtCa^2+^ extrusion and resulting in mtCa^2+^ overload, enhances HIF1α levels through an increase in mtROS. Our xenograft model shows that despite the reduced tumor burden and tumor size in SCID mice injected with NCLX KO HCT116 cells compared to SCID mice injected with control HCT116 cells, the survival of the former mice was significantly reduced compared to the latter. This result can be explained by the enhanced metastasis to the liver and colon in SCID mice injected with NCLX KO HCT116 cells. Indeed, we were able to show that the increased invasive properties of NCLX KO CRC cells were associated with increased MMP1, MMP2, and MMP9 activities, mRNA and protein levels, which are known to be regulated by HIF1α (Ben-Yosef et al., 2002; Choudhry and Harris, 2018; Muñoz-Nájar et al., 2006; Shin et al., 2015; Tsai et al., 2016). We showed that NCLX KO CRC cells are chemoresistant to treatment by 5-FU, with both proliferation and migration of NCLX KO CRC cells not significantly altered by 5-FU treatment. It is feasible to suspect that the decrease in proliferation contributes to the chemoresistance phenotype of cells lacking NCLX. In previous work, we demonstrated that decreased mtCa^2+^ extrusion also leads to a decrease in cytosolic Ca^2+^ and reduced store-operated Ca^2+^ entry (SOCE) mediated by ORAI1 channels (Ben-Kasus Nissim et al., 2017). Similarly, we see a decrease in cytosolic Ca^2+^ and SOCE when NCLX is knocked out in CRC cell lines (**Figure S4O, P**). A consequence of reducing SOCE and cytosolic Ca^2+^ is decreased activation of the Ca^2+^ sensitive transcription factor NFAT (Trebak and Kinet, 2019). NFAT is a known stimulator of MYC activation (Buchholz et al., 2006; Mognol et al., 2012; Singh et al., 2010), suggesting that reduced NFAT signaling as a consequence of NCLX downregulation might be contributing to the decrease in Myc pathway activation, cell cycle progression, and proliferation (**Figures 6A, B, D, Figure S6D**).

The EMT phenotype and decreases in proliferation are hallmarks of cancer stem cells and the presence of cancer stem cells within tumors is one of the major causes of chemotherapy resistance (Izumiya et al., 2012; Munro et al., 2018; Ohata et al., 2019; Zhao, 2016). Studies have shown that CSCs are enriched on treatment with 5-FU through different molecular mechanisms (Abdullah and Chow, 2013; Lu et al., 2015; Touil et al., 2014). Recent work showed that increased chemoresistance of breast cancer was mediated by HIF1α-mediated glutathione biosynthesis and glutathione-mediated enrichment with cancer stem cells (Lu et al., 2015). Chemotherapy in breast cancer induces HIF1α-dependent glutathione biosynthesis by enhancing the expression of the cystine transporter xCT (SLC7A11) and the regulatory subunit of glutamate-cysteine ligase (GCLM) (Lu et al., 2015). The glutathione thus generated inhibits the MEK/ERK pathway through copper chelation resulting in enhanced expression of the stem cell markers NANOG, Oct4 and Sox2. We similarly observed enhanced expression of SLC7A11 and GCLM in NCLX KO clones from both HCT116 and DLD1 cells (**Figure S7A-I**). In addition to contributing to stem cell regulation, an increase in enzymes responsible for glutathione synthesis potentially provides additional survival advantages to NCLX knockout cells, by enhancing the ROS scavenging ability of CRC cells and preventing any lethal build-up of ROS as a consequence of mtCa^2+^ elevation.

In summary, we have identified a novel signaling mitochondrial Ca^2+^/ROS signaling axis in colorectal cancer that is initiated in response to decreased NCLX expression, as commonly observed in CRC patient tumors. While reduced mtCa^2+^ extrusion causes mitochondrial depolarization, mitochondrial dysfunction, reduced cell cycle-related gene expression, increased mitophagy and reduced proliferation, leading to smaller tumors both in the colitis-associated CRC model and the xenograft CRC model, the mtCa^2+^ overload induced by NCLX loss enhances mtROS, which in turn enhances CRC metastasis through HIF1α-dependent increases in glycolysis, chemoresistance and pro-metastatic gene expression signatures, contributing to increased metastatic burden and enhanced lethality of SCID mice injected with NCLX KO CRC cells. These changes are reminiscent of the highly metastatic mesenchymal CRC subtype, and our work suggests that restoring mtCa^2+^ homeostasis in CRC tumors might be beneficial in limiting or preventing CRC progression and metastasis.

## Acknowledgments

The authors acknowledge Dr. Han Chen from the Pennsylvania State University College of Medicine EM facility for assistance with TEM imaging and Dr. Katherine M. Aird (The Pennsylvania State University) for providing shRNA of NCLX. Our study was supported by the National Heart, Lung, and Blood Institute (R01-HL123364, R01-HL097111, and R35-HL150778 to M.T.), National Institute on Aging (R21-AG050072 to M.T.), and American Heart Association Postdoctoral Fellowship (9POST34380606) to T.P. G.S.Y and W.A.K are supported by the Peter and Marshia Carlino Fund for IBD research. We also acknowledge, Flow Cytometry & Cell Sorting, Imaging, Informatics & Data Analysis Core facility (The Pennsylvania State University, College of Medicine).

## Author Contributions

Conceptualization, T.P., M.G., N.H., and M.T.; Methodology, T.P. and M.T.; Validation, C.D., and R.Y.; Formal analysis, T.P., V.W., and C.D.; Investigation, T.P., C.D., M.G., M.J., X.Z., and S.E.; Resources, T.P., P.X., G.S.Y., W.A.K., I.S., and V.W.; Writing, T.P., and M.T. with inputs from all authors; Supervision, M.T.; Funding Acquisition, M.T.

## References

Abdullah, L. N., and Chow, E. K.-H. (2013). Mechanisms of chemoresistance in cancer stem cells. Clin Transl Med 2, 3–3.

Ashrafi, G., and Schwarz, T. L. (2013). The pathways of mitophagy for quality control and clearance of mitochondria. Cell Death & Differentiation 20, 31–42.

Baughman, J. M., Perocchi, F., Girgis, H. S., Plovanich, M., Belcher-Timme, C. A., Sancak, Y., Bao, X. R., Strittmatter, L., Goldberger, O., Bogorad, R. L., et al. (2011). Integrative genomics identifies MCU as an essential component of the mitochondrial calcium uniporter. Nature 476, 341–345.

Bell, E. L., Klimova, T. A., Eisenbart, J., Schumacker, P. T., and Chandel, N. S. (2007). Mitochondrial reactive oxygen species trigger hypoxia-inducible factor-dependent extension of the replicative life span during hypoxia. Molecular and cellular biology 27, 5737–5745.

Bellot, G., Garcia-Medina, R., Gounon, P., Chiche, J., Roux, D., Pouysségur, J., and Mazure, N. M. (2009). Hypoxia-Induced Autophagy Is Mediated through Hypoxia-Inducible Factor Induction of BNIP3 and BNIP3L via Their BH3 Domains. Molecular and cellular biology 29, 2570–2581.

Ben-Kasus Nissim, T., Zhang, X., Elazar, A., Roy, S., Stolwijk, J. A., Zhou, Y., Motiani, R. K., Gueguinou, M., Hempel, N., Hershfinkel, M., et al. (2017). Mitochondria control store-operated Ca(2+) entry through Na(+) and redox signals. Embo j 36, 797–815.

Ben-Yosef, Y., Lahat, N., Shapiro, S., Bitterman, H., and Miller, A. (2002). Regulation of endothelial matrix metalloproteinase-2 by hypoxia/reoxygenation. Circulation research 90, 784–791.

Bernardi, P., and Di Lisa, F. (2015). The mitochondrial permeability transition pore: molecular nature and role as a target in cardioprotection. Journal of molecular and cellular cardiology 78, 100–106.

Bertero, E., and Maack, C. (2018). Calcium Signaling and Reactive Oxygen Species in Mitochondria. Circulation Research 122, 1460–1478.

Blank, A., Roberts, D. E., 2nd, Dawson, H., Zlobec, I., and Lugli, A. (2018). Tumor Heterogeneity in Primary Colorectal Cancer and Corresponding Metastases. Does the Apple Fall Far From the Tree? Front Med (Lausanne) 5, 234–234.

Booth, D. M., Enyedi, B., Geiszt, M., Varnai, P., and Hajnoczky, G. (2016). Redox Nanodomains Are Induced by and Control Calcium Signaling at the ER-Mitochondrial Interface. Mol Cell 63, 240–248.

Brookes, P. S., Yoon, Y., Robotham, J. L., Anders, M. W., and Sheu, S. S. (2004). Calcium, ATP, and ROS: a mitochondrial love-hate triangle. Am J Physiol Cell Physiol 287, C817–833.

Bu, P., Chen, K. Y., Xiang, K., Johnson, C., Crown, S. B., Rakhilin, N., Ai, Y., Wang, L., Xi, R., Astapova, I., et al. (2018). Aldolase B-Mediated Fructose Metabolism Drives Metabolic Reprogramming of Colon Cancer Liver Metastasis. Cell Metab 27, 1249–1262e1244.

Buchholz, M., Schatz, A., Wagner, M., Michl, P., Linhart, T., Adler, G., Gress, T. M., and Ellenrieder, V. (2006). Overexpression of c-myc in pancreatic cancer caused by ectopic activation of NFATc1 and the Ca2+/calcineurin signaling pathway. The EMBO journal 25, 3714–3724.

Burns, J. S., and Manda, G. (2017). Metabolic Pathways of the Warburg Effect in Health and Disease: Perspectives of Choice, Chain or Chance. Int J Mol Sci 18.

Buvinic, S., Almarza, G., Bustamante, M., Casas, M., López, J., Riquelme, M., Sáez, J. C., Huidobro-Toro, J. P., and Jaimovich, E. (2009). ATP released by electrical stimuli elicits calcium transients and gene expression in skeletal muscle. J Biol Chem 284, 34490–34505.

Celsi, F., Pizzo, P., Brini, M., Leo, S., Fotino, C., Pinton, P., and Rizzuto, R. (2009). Mitochondria, calcium and cell death: A deadly triad in neurodegeneration. Biochimica et Biophysica Acta (BBA) - Bioenergetics 1787, 335–344.

Chandrashekar, D. S., Bashel, B., Balasubramanya, S. A. H., Creighton, C. J., Ponce-Rodriguez, I., Chakravarthi, B. V. S. K., and Varambally, S. (2017). UALCAN: A Portal for Facilitating Tumor Subgroup Gene Expression and Survival Analyses. Neoplasia 19, 649–658.

Choudhry, H., and Harris, A. L. (2018). Advances in Hypoxia-Inducible Factor Biology. Cell Metab 27, 281–298.

Chourasia, A. H., Tracy, K., Frankenberger, C., Boland, M. L., Sharifi, M. N., Drake, L. E., Sachleben, J. R., Asara, J. M., Locasale, J. W., Karczmar, G. S., and Macleod, K. F. (2015). Mitophagy defects arising from BNip3 loss promote mammary tumor progression to metastasis. EMBO reports 16, 1145–1163.

Dan Dunn, J., Alvarez, L. A. J., Zhang, X., and Soldati, T. (2015). Reactive oxygen species and mitochondria: A nexus of cellular homeostasis. Redox Biology 6, 472–485.

Danese, A., Patergnani, S., Bonora, M., Wieckowski, M. R., Previati, M., Giorgi, C., and Pinton, P. (2017). Calcium regulates cell death in cancer: Roles of the mitochondria and mitochondria-associated membranes (MAMs). Biochimica et Biophysica Acta (BBA) - Bioenergetics 1858, 615–627.

De Stefani, D., Raffaello, A., Teardo, E., Szabò, I., and Rizzuto, R. (2011). A forty-kilodalton protein of the inner membrane is the mitochondrial calcium uniporter. Nature 476, 336–340.

De Stefani, D., Rizzuto, R., and Pozzan, T. (2016). Enjoy the Trip: Calcium in Mitochondria Back and Forth. Annual Review of Biochemistry 85, 161–192.

Dodd, K. M., Yang, J., Shen, M. H., Sampson, J. R., and Tee, A. R. (2015). mTORC1 drives HIF-1α and VEGF-A signalling via multiple mechanisms involving 4E-BP1, S6K1 and STAT3. Oncogene 34, 2239–2250.

Dunne, P. D., Alderdice, M., O’Reilly, P. G., Roddy, A. C., McCorry, A. M. B., Richman, S., Maughan, T., McDade, S. S., Johnston, P. G., Longley, D. B., et al. (2017). Cancer-cell intrinsic gene expression signatures overcome intratumoural heterogeneity bias in colorectal cancer patient classification. Nature communications 8, 15657–15657.

Fan, P., Xie, X. H., Chen, C. H., Peng, X., Zhang, P., Yang, C., and Wang, Y. T. (2019). Molecular Regulation Mechanisms and Interactions Between Reactive Oxygen Species and Mitophagy. DNA Cell Biol 38, 10–22.

Frank, M., Duvezin-Caubet, S., Koob, S., Occhipinti, A., Jagasia, R., Petcherski, A., Ruonala, M. O., Priault, M., Salin, B., and Reichert, A. S. (2012). Mitophagy is triggered by mild oxidative stress in a mitochondrial fission dependent manner. Biochimica et Biophysica Acta (BBA) - Molecular Cell Research 1823, 2297–2310.

Gillies, R. J., Robey, I., and Gatenby, R. A. (2008). Causes and Consequences of Increased Glucose Metabolism of Cancers. Journal of Nuclear Medicine 49, 24S–42S.

Giorgi, C., Baldassari, F., Bononi, A., Bonora, M., De Marchi, E., Marchi, S., Missiroli, S., Patergnani, S., Rimessi, A., Suski, J. M., et al. (2012). Mitochondrial Ca(2+) and apoptosis. Cell calcium 52, 36–43.

Gonzalez, F. A., Rozengurt, E., and Heppel, L. A. (1989). Extracellular ATP induces the release of calcium from intracellular stores without the activation of protein kinase C in Swiss 3T6 mouse fibroblasts. Proceedings of the National Academy of Sciences of the United States of America 86, 4530–4534.

Guinney, J., Dienstmann, R., Wang, X., de Reyniès, A., Schlicker, A., Soneson, C., Marisa, L., Roepman, P., Nyamundanda, G., Angelino, P., et al. (2015). The consensus molecular subtypes of colorectal cancer. Nature medicine 21, 1350–1356.

Halestrap, A. P. (2009). What is the mitochondrial permeability transition pore? Journal of Molecular and Cellular Cardiology 46, 821–831.

Hamanaka, R. B., and Chandel, N. S. (2010). Mitochondrial reactive oxygen species regulate cellular signaling and dictate biological outcomes. Trends Biochem Sci 35, 505–513.

Han, T., Kang, D., Ji, D., Wang, X., Zhan, W., Fu, M., Xin, H.-B., and Wang, J.-B. (2013). How does cancer cell metabolism affect tumor migration and invasion? Cell Adh Migr 7, 395–403.

Hansford, R. G. (1994). Physiological role of mitochondrial Ca2+ transport. J Bioenerg Biomembr 26, 495–508.

Hempel, N., and Trebak, M. (2017). Crosstalk between calcium and reactive oxygen species signaling in cancer. Cell Calcium 63, 70–96.

Hudson, C. C., Liu, M., Chiang, G. G., Otterness, D. M., Loomis, D. C., Kaper, F., Giaccia, A. J., and Abraham, R. T. (2002). Regulation of Hypoxia-Inducible Factor 1α Expression and Function by the Mammalian Target of Rapamycin. Molecular and cellular biology 22, 7004–7014.

Isella, C., Brundu, F., Bellomo, S. E., Galimi, F., Zanella, E., Porporato, R., Petti, C., Fiori, A., Orzan, F., Senetta, R., et al. (2017). Selective analysis of cancer-cell intrinsic transcriptional traits defines novel clinically relevant subtypes of colorectal cancer. Nature communications 8, 15107–15107.

Izumiya, M., Kabashima, A., Higuchi, H., Igarashi, T., Sakai, G. E. N., Iizuka, H., Nakamura, S., Adachi, M., Hamamoto, Y., Funakoshi, S., et al. (2012). Chemoresistance Is Associated with Cancer Stem Cell-like Properties and Epithelial-to-Mesenchymal Transition in Pancreatic Cancer Cells. Anticancer Research 32, 3847–3853.

Jin, S. M., Lazarou, M., Wang, C., Kane, L. A., Narendra, D. P., and Youle, R. J. (2010). Mitochondrial membrane potential regulates PINK1 import and proteolytic destabilization by PARL. The Journal of cell biology 191, 933–942.

Kerkhofs, M., Bittremieux, M., Morciano, G., Giorgi, C., Pinton, P., Parys, J. B., and Bultynck, G. (2018). Emerging molecular mechanisms in chemotherapy: Ca2+ signaling at the mitochondria-associated endoplasmic reticulum membranes. Cell Death & Disease 9, 334.

Koval, O. M., Nguyen, E. K., Santhana, V., Fidler, T. P., Sebag, S. C., Rasmussen, T. P., Mittauer, D. J., Strack, S., Goswami, P. C., Abel, E. D., and Grumbach, I. M. (2019). Loss of MCU prevents mitochondrial fusion in G1-S phase and blocks cell cycle progression and proliferation. Sci Signal 12.

Lehmann, S., te Boekhorst, V., Odenthal, J., Bianchi, R., van Helvert, S., Ikenberg, K., Ilina, O., Stoma, S., Xandry, J., Jiang, L., et al. (2017). Hypoxia Induces a HIF-1-Dependent Transition from Collective-to-Amoeboid Dissemination in Epithelial Cancer Cells. Current Biology 27, 392–400.

Liberti, M. V., and Locasale, J. W. (2016). The Warburg Effect: How Does it Benefit Cancer Cells? Trends Biochem Sci 41, 211–218.

Lu, H., Samanta, D., Xiang, L., Zhang, H., Hu, H., Chen, I., Bullen, J. W., and Semenza, G. L. (2015). Chemotherapy triggers HIF-1–dependent glutathione synthesis and copper chelation that induces the breast cancer stem cell phenotype. Proceedings of the National Academy of Sciences 112, E4600–E4609.

Luongo, T. S., Lambert, J. P., Gross, P., Nwokedi, M., Lombardi, A. A., Shanmughapriya, S., Carpenter, A. C., Kolmetzky, D., Gao, E., van Berlo, J. H., et al. (2017). The mitochondrial Na(+)/Ca(2+) exchanger is essential for Ca(2+) homeostasis and viability. Nature 545, 93–97.

Marchi, S., Lupini, L., Patergnani, S., Rimessi, A., Missiroli, S., Bonora, M., Bononi, A., Corra, F., Giorgi, C., De Marchi, E., et al. (2013). Downregulation of the mitochondrial calcium uniporter by cancer-related miR-25. Curr Biol 23, 58–63.

Masoud, G. N., and Li, W. (2015). HIF-1α pathway: role, regulation and intervention for cancer therapy. Acta Pharm Sin B 5, 378–389.

McCormack, J. G., Halestrap, A. P., and Denton, R. M. (1990). Role of calcium ions in regulation of mammalian intramitochondrial metabolism. Physiol Rev 70, 391–425.

Mognol, G. P., de Araujo-Souza, P. S., Robbs, B. K., Teixeira, L. K., and Viola, J. P. B. (2012). Transcriptional regulation of the c-Myc promoter by NFAT1 involves negative and positive NFAT-responsive elements. Cell Cycle 11, 1014–1028.

Montero, M., Alonso, M. T., Carnicero, E., Cuchillo-Ibanez, I., Albillos, A., Garcia, A. G., Garcia-Sancho, J., and Alvarez, J. (2000). Chromaffin-cell stimulation triggers fast millimolar mitochondrial Ca2+ transients that modulate secretion. Nat Cell Biol 2, 57–61.

Muñoz-Nájar, U. M., Neurath, K. M., Vumbaca, F., and Claffey, K. P. (2006). Hypoxia stimulates breast carcinoma cell invasion through MT1-MMP and MMP-2 activation. Oncogene 25, 2379–2392.

Munro, M. J., Wickremesekera, S. K., Peng, L., Tan, S. T., and Itinteang, T. (2018). Cancer stem cells in colorectal cancer: a review. Journal of Clinical Pathology 71, 110.

Ohata, H., Shiokawa, D., Obata, Y., Sato, A., Sakai, H., Fukami, M., Hara, W., Taniguchi, H., Ono, M., Nakagama, H., and Okamoto, K. (2019). NOX1-Dependent mTORC1 Activation via S100A9 Oxidation in Cancer Stem-like Cells Leads to Colon Cancer Progression. Cell Reports 28, 1282–1295.e1288.

Palty, R., Silverman, W. F., Hershfinkel, M., Caporale, T., Sensi, S. L., Parnis, J., Nolte, C., Fishman, D., Shoshan-Barmatz, V., Herrmann, S., et al. (2010). NCLX is an essential component of mitochondrial Na+/Ca2+ exchange. Proc Natl Acad Sci U S A 107, 436–441.

Pan, Y., Mansfield, K. D., Bertozzi, C. C., Rudenko, V., Chan, D. A., Giaccia, A. J., and Simon, M. C. (2007). Multiple factors affecting cellular redox status and energy metabolism modulate hypoxia-inducible factor prolyl hydroxylase activity in vivo and in vitro. Molecular and cellular biology 27, 912–925.

Pathak, T., and Trebak, M. (2018). Mitochondrial Ca(2+) signaling. Pharmacol Ther 192, 112–123.

Paupe, V., and Prudent, J. (2018). New insights into the role of mitochondrial calcium homeostasis in cell migration. Biochemical and Biophysical Research Communications 500, 75–86.

Pinton, P., Giorgi, C., Siviero, R., Zecchini, E., and Rizzuto, R. (2008). Calcium and apoptosis: ER-mitochondria Ca2+ transfer in the control of apoptosis. Oncogene 27, 6407–6418.

Polyak, K., and Weinberg, R. A. (2009). Transitions between epithelial and mesenchymal states: acquisition of malignant and stem cell traits. Nature Reviews Cancer 9, 265–273.

Porporato, P. E., Filigheddu, N., Pedro, J. M. B.-S., Kroemer, G., and Galluzzi, L. (2018). Mitochondrial metabolism and cancer. Cell Research 28, 265–280.

Ren, T., Zhang, H., Wang, J., Zhu, J., Jin, M., Wu, Y., Guo, X., Ji, L., Huang, Q., Zhang, H., et al. (2017). MCU-dependent mitochondrial Ca(2+) inhibits NAD(+)/SIRT3/SOD2 pathway to promote ROS production and metastasis of HCC cells. Oncogene 36, 5897–5909.

Reya, T., Morrison, S. J., Clarke, M. F., and Weissman, I. L. (2001). Stem cells, cancer, and cancer stem cells. Nature 414, 105–111.

Santulli, G., Xie, W., Reiken, S. R., and Marks, A. R. (2015). Mitochondrial calcium overload is a key determinant in heart failure. Proceedings of the National Academy of Sciences 112, 11389–11394.

Schofield, J. H., and Schafer, Z. T. (2020). Mitochondrial ROS and Mitophagy: A Complex and Nuanced Relationship. Antioxid Redox Signal.

Semenza, G. L. (2003). Targeting HIF-1 for cancer therapy. Nature Reviews Cancer 3, 721–732.

Shin, D. H., Dier, U., Melendez, J. A., and Hempel, N. (2015). Regulation of MMP-1 expression in response to hypoxia is dependent on the intracellular redox status of metastatic bladder cancer cells. Biochimica et biophysica acta 1852, 2593–2602.

Siegel, R. L., Miller, K. D., and Jemal, A. (2020). Cancer statistics, 2020. CA: A Cancer Journal for Clinicians 70, 7–30.

Singh, G., Singh, S. K., König, A., Reutlinger, K., Nye, M. D., Adhikary, T., Eilers, M., Gress, T. M., Fernandez-Zapico, M. E., and Ellenrieder, V. (2010). Sequential activation of NFAT and c-Myc transcription factors mediates the TGF-beta switch from a suppressor to a promoter of cancer cell proliferation. The Journal of biological chemistry 285, 27241–27250.

Starkov, A. A. (2008). The role of mitochondria in reactive oxygen species metabolism and signaling. Ann N Y Acad Sci 1147, 37–52.

Tandon, P., Gallo, C. A., Khatri, S., Barger, J. F., Yepiskoposyan, H., and Plas, D. R. (2011). Requirement for ribosomal protein S6 kinase 1 to mediate glycolysis and apoptosis resistance induced by Pten deficiency. Proceedings of the National Academy of Sciences 108, 2361–2365.

Tennant, D. A., Durán, R. V., Boulahbel, H., and Gottlieb, E. (2009). Metabolic transformation in cancer. Carcinogenesis 30, 1269–1280.

Tennant, D. A., Durán, R. V., and Gottlieb, E. (2010). Targeting metabolic transformation for cancer therapy. Nature reviews Cancer 10, 267–277.

Tosatto, A., Sommaggio, R., Kummerow, C., Bentham, R. B., Blacker, T. S., Berecz, T., Duchen, M. R., Rosato, A., Bogeski, I., Szabadkai, G., et al. (2016). The mitochondrial calcium uniporter regulates breast cancer progression via HIF-1alpha. EMBO Mol Med 8, 569–585.

Touil, Y., Igoudjil, W., Corvaisier, M., Dessein, A. F., Vandomme, J., Monte, D., Stechly, L., Skrypek, N., Langlois, C., Grard, G., et al. (2014). Colon cancer cells escape 5FU chemotherapy-induced cell death by entering stemness and quiescence associated with the c-Yes/YAP axis. Clin Cancer Res 20, 837–846.

Trebak, M., and Kinet, J.-P. (2019). Calcium signalling in T cells. Nature Reviews Immunology 19, 154–169.

Tsai, S.-H., Huang, P.-H., Hsu, Y.-J., Peng, Y.-J., Lee, C.-H., Wang, J.-C., Chen, J.-W., and Lin, S.-J. (2016). Inhibition of hypoxia inducible factor-1α attenuates abdominal aortic aneurysm progression through the down-regulation of matrix metalloproteinases. Scientific Reports 6, 28612.

Twig, G., and Shirihai, O. S. (2011). The interplay between mitochondrial dynamics and mitophagy. Antioxid Redox Signal 14, 1939–1951.

Vultur, A., Gibhardt, C. S., Stanisz, H., and Bogeski, I. (2018). The role of the mitochondrial calcium uniporter (MCU) complex in cancer. Pflugers Arch 470, 1149–1163.

Vyas, S., Zaganjor, E., and Haigis, M. C. (2016). Mitochondria and Cancer. Cell 166, 555–566.

Wigerup, C., Påhlman, S., and Bexell, D. (2016). Therapeutic targeting of hypoxia and hypoxia-inducible factors in cancer. Pharmacology & Therapeutics 164, 152–169.

Wu, H., Carvalho, P., and Voeltz, G. K. (2018). Here, there, and everywhere: The importance of ER membrane contact sites. Science 361, eaan5835.

Youle, R. J., and Narendra, D. P. (2010). Mechanisms of mitophagy. Nature Reviews Molecular Cell Biology 12, 9.

Zhang, H., Bosch-Marce, M., Shimoda, L. A., Tan, Y. S., Baek, J. H., Wesley, J. B., Gonzalez, F. J., and Semenza, G. L. (2008). Mitochondrial autophagy is an HIF-1-dependent adaptive metabolic response to hypoxia. The Journal of biological chemistry 283, 10892–10903.

Zhang, J., and Ney, P. A. (2009). Role of BNIP3 and NIX in cell death, autophagy, and mitophagy. Cell Death & Differentiation 16, 939–946.

Zhao, J. (2016). Cancer stem cells and chemoresistance: The smartest survives the raid. Pharmacology & Therapeutics 160, 145–158.

