## Supplemental file for "Colorectal adenocarcinomas downregulate the mitochondrial Na^+^/Ca^2+^ exchanger NCLX to drive metastatic spread"

**Pathak et al.**

**Online Supplement**

**Methods**
**Key resource table**

| **Reagent or Resources** | **Source** | **Identifier** |
|---|---|---|
| Antibodies | | |
| Hif1α | Cell Signaling Technology | Cat# 14179s |
| ALDOA | Cell Signaling Technology | Cat# 8060S |
| HK2 | Cell Signaling Technology | Cat# 2867S |
| LDHA | Cell Signaling Technology | Cat# 2012S |
| MMP1 | Abcam | Cat# ab38929 |
| MMP2 | Santa Cruz Biotechnology | Cat# sc-13594 |
| MMP9 | Abcam | Cat# ab73734 |
| LC3B | Abcam | Cat# ab51520 |
| OXPHOS | Abcam | Cat# ab110413 |
| GAPDH | Millipore Sigma | Cat# MAB374 |
| αTubulin | Cell Signaling Technology | Cat# 3873S |
| HSC70 | Santa Cruz Biotechnology | Cat# sc-24 |
| p62 | Abcam | Cat# ab109012 |
| Cleaved Caspase3 | Cell Signaling Technology | Cat# 9661S |
| pAMPK | Cell Signaling Technology | Cat# 2535S |
| AMPK | Cell Signaling Technology | Cat# 5831S |
| pS6K | Cell Signaling Technology | Cat# 9234S |
| S6K | Cell Signaling Technology | Cat# 2708S |
| IRDye 800CW Goat anti-Mouse | Li-Core Biosciences | Cat# 925-32210 |
| IRDye 800CW Donkey anti-Rabbit | Li-Core Biosciences | Cat# 925-32213 |
| IRDye 680RD Goat anti-Mouse | Li-Core Biosciences | Cat# 925-68070 |
| IRDye 680RD Goat anti-Rabbit | Li-Core Biosciences | Cat# 925-68071 |
| **Chemicals** | | |
| **Chemical Name** | **Source** | **Identifier** |
| DAPI | Sigma-Aldrich | Cat# D9542 |
| Hoechst | Thermo Fisher Scientific | Cat# H3570 |
| MitoSox Red | Thermo Fisher Scientific | Cat# M36008 |

| | | |
|---|---|---|
| Mito TEMPO | Thermo Fisher Scientific | Cat# SML0737 |
| Mito Tracker Green FM | Thermo Fisher Scientific | Cat# M7514 |
| Mito Tracker Deep red FM | Cell Signaling Technology | Cat# 8778S |
| Fura-2 AM | Thermo Fisher Scientific | Cat# F1221 |
| FBS | VWR | Cat# MISC-LFSCI-STP |
| Antibiotic and Antimycotic | Thermo Fisher Scientific | Cat# 15240062 |
| McCoy's 5A | Corning | Cat# 10-050CV |
| RPMI-1640 | Corning | Cat# 10-040CV |
| Lipofectamine 2000 | Thermo Fisher Scientific | Cat# 11668019 |
| TrypLE | Thermo Fisher Scientific | Cat# 12605028 |
| $CoCl_2$ | Sigma-Aldrich | Cat# 15862 |
| 2-deoxy-D-glucose (2-DG) | Sigma-Aldrich | Cat# D8375 |
| 5- Fluorouracil | Sigma-Aldrich | Cat# F6627 |
| Glucose | Sigma-Aldrich | Cat# D9434 |
| Puromycin | MP Biomedical | Cat# 02100552 |
| RIPA buffer | Sigma | Cat# R0278 |
| ATP | Sigma | Cat# A9187 |
| Dextrose | Fisher Scientific | Cat# D14 |
| Tris Base | Fisher Scientific | Cat# BP152-5 |
| NaCl | Fisher Scientific | Cat# S671 |
| MOPS SDS running buffer | Thermo Fisher Scientific | Cat# NP0001 |
| Tris-Glycine transfer buffer | Bio-rad | Cat#161-0734 |
| KCl | Fisher Scientific | Cat# P217 |
| $MgCl_2$ | Fisher Scientific | Cat# M33 |
| $CaCl_2$ | Fisher Scientific | Cat# C614 |
| HEPES | Fisher Scientific | Cat# BP310 |
| LDS sample buffer | Thermo Fisher Scientific | Cat# NP0007 |
| NuPAGE Bis-Tris precast gels | Thermo Fisher Scientific | Cat# NP0321 |
| Polyvinylidene difluoride membrane | Li-Core Biosciences | Cat# 88518 |
| Odyssey Blocking Buffer (TBS) | Li-Core Biosciences | Cat# 937-50003 |
| Dextran sulfate sodium | MP Biomedical | Cat# 0216011080 |
| Azoxymethane | Sigma | Cat# A5486 |
| FluoroBlok | Corning | Cat# 351152 |
| BioCoat™ Tumor Invasion Plate | Corning | Cat# 80774380 |
| BCA assay kit | Thermo Fisher Scientific | Cat# A53225 |
| TMRE | Thermo Fisher Scientific | Cat# T669 |
| CyQUANT | Thermo Fisher Scientific | Cat# C35006 |
| cDNA Reverse Transcription Kit | Applied biosystems | Cat# 4368814 |
| DNase I | Thermo Fisher Scientific | Cat# 18068-015 |
| TRIzol | Thermo Fisher Scientific | Cat# 15596018 |
| SYBER select master mix | Thermo Fisher Scientific | Cat# 4472920 |
| Seahorse XF DMEM Medium pH 7.4 | Agilent Technologies | Cat# 103575-100 |

| | | |
|---|---|---|
| Seahorse XF 100 mM pyruvate solution | Agilent Technologies | Cat# 103578-100 |
| Seahorse XF 200 mM glutamine solution | Agilent Technologies | Cat# 103579-100 |
| Seahorse XF 1.0 M glucose solution | Agilent Technologies | Cat# 103577-100 |
| Seahorse XFp Mito Fuel Flex Test Kit | Agilent Technologies | Cat# 103270-100 |
| Seahorse XFp Glycolysis Stress Test Kit | Agilent Technologies | Cat# 103017-100 |
| Seahorse XFp Cell Mito Stress Test Kit | Agilent Technologies | Cat# 103010-100 |
| Zymogram Developing Buffer (10X) | Thermo Fisher Scientific | Cat# LC2671 |
| Zymogram Renaturing Buffer (10X) | Thermo Fisher Scientific | Cat# LC2670 |
| Tris-Glycine SDS Running Buffer (10X) | Thermo Fisher Scientific | Cat# LC2675 |
| Novex™ 10% Zymogram Plus (Gelatin) Protein Gels, 1.0 mm, 10-well | Thermo Fisher Scientific | Cat# ZY00100 |
| SimplyBlue™ Safe Stain | Thermo Fisher Scientific | Cat# LC6060 |
| Tween 20 | Fisher Scientific | Cat# BP337 |
| **Cell Lines** | | |
| **Cell line Name** | **Source** | **Identifier** |
| Human colon cancer HCT116 cells (male) | ATCC | ATCC# CCL-247 |
| Human colon cancer HT29 cells (female) | ATCC | ATCC# HTB-38 |
| Human colon cancer DLD1 cells (male) | ATCC | ATCC# CCL-221 |
| Human colon cancer HCT116 NCLX KO cells | This paper | N/A |
| Human colon cancer DLD1 NCLX KO cells | This paper | N/A |
| Human colon cancer HCT116 shNCLX cells | This paper | N/A |
| Human colon cancer DLD1 shNCLX cells | This paper | N/A |
| **RT-qPCR primer sequences** | | |
| **Gene name** | **Forward Primer** | **Reverse Primer** |
| SLC7A11 | AGGGTCACCTTCCAGAAATC | GAAGATAAATCAGCCCAGCA |

| | | |
|---|---|---|
| GCLM | CATTTACAGCCTTAC TGGGAGG | ATGCAGTCAAATCTGGT GGCA |
| FOX3 | CGGCTTCGGCTCTTA GCAAA | CGGACAAACGGCTCACT CT |
| NANOG | TTTGTGGGCCTGAAG AAAACT | AGGGCTGTCCTGAATAA GCAG |
| OCT4 | TTCAGCCAAACGACC ATCTG | CACGAGGGTTTCTGCTT TGC |
| SOX2 | GCCGAGTGGAAACTT TTGTCG | GGCAGCGTGTACTTATC CTTCT |
| GLUT1 | TATCGTCAACACGGC CTTCACT | AACAGCTCCTCGGGTGT CTTAT |
| HK2 | GCCATCCTGCAACAC TTAGGG | GTGAGGATGTAGCTTGT AGAGGGT |
| GPI | TGTGTTCACCAAGCT CACAC | GTAGAAGCGTCGTGAG AGGT |
| ALDOA | AGGCCATGCTTGCAC TCAG | AGGGCCCAGGGCTTCAG |
| ENO1 | GACTTGGCTGGCAAC TCTG | GGTCATCGGGAGACTTG AAG |
| LDHA | GGTTGGTGCTGTTGG CATGG | TGCCCCAGCCGTGATAA TGA |
| MMP1 | ATGCTGAAACCCTGA AGGTG | GAGCATCCCCTCCAATA CCT |
| MMP2 | ACCAGCTGGCCTAGT GATGATG | GGCTTCCGCATGGTCTC GATG |
| MMP9 | ACGCACGACGTCTTC CAGTA | CCACCTGGTTCAACTCA CTCC |
| qNCLX_1 | GCCAGCATTTGTGTC CATTT | AATTCGTCTCGGCCACT TAC |
| qNCLX_2 | CCGGCAGAAGGCTG AATCTG | ACCTTGCGGCAGTCTAC CAC |
| MDR1 | AACGGAAGCCAGAA CATTCC | AGGCTTCCTGTGGCAAA GAG |
| Tubulin | AGTCCAAGCTGGAGT TCTCTAT | CAATCAGAGTGCTCCAG GGT |
| GAPDH | GGGAAACCCATCAC CATCTT | CCAGTAGACTCCACGAC ATACT |
| G6PD | CGAGGCCGTCACCA AGAAC | GTAGTGGTCGATGCGGT AGA |
| PGD | ATGGCCCAAGCTGAC ATCG | AAAGCCGTGGTCATTCA TGTT |
| TKT | TCCACACCATGCGCT ACAAG | CAAGTCGGAGCTGATCT TCCT |
| **Guide RNA Sequences** | | |

| Name | Sequence | Figure # |
|---|---|---|
| g1 | GCGCAGATTCAGCCT TCTGC | Figure 2A and S3A |
| g2 | GGGATACTCACGTCT ACCAC | Figure S3B and C |
| g3 | GTAGACGTGAGTATC CCGGT | Figure S3E |
| g4 | ACCCACACCAGCAGT CCGTC | Figure S3E |
| **shRNA and siRNA sequences** | | |
| **Sequence name** | **Sequence** | **Figure #** |
| shNCLX#2 | GCCTTCTTGCTGTCA TGCAAT | Figure S3I |
| shNCLX#3 | GCTCCTCTTCTACCT GAACTT | Figure S3I |
| siNCLX | AACGGCCCCUCAAC UGUCUT | Figure S4E |
| **Primers for screening genomic DNA of NCLX KO clones** | | |
| **Name** | **Sequence** | **Figure #** |
| NCLX_1 | GCGTGCTGGTTACCA CAG T | Figure S3F |
| NCLX_2 | CCACGGAAGAGCAT GAGGAA | Figure S3F |
| NCLX_3 | ACTTAGCACATCGCC ACC TG | Figure S3F |
| NCLX_4 | CTGATCTGCACGCTG AAT GG | Figure S3F |
| NCLX_5 | GAGGTACACAGCAG TTCT CCC | Figure S3F |
| NCLX_6 | CAGCTGGTGCCCTCA AAC AC | Figure S3F |
| **Primers for genotyping NCLX$^{-/-}$ mice** | | |
| **Name** | **Sequence** | **Figure #** |
| px77 | TACAGTCTGGCTCGT TCC CT | Figure S1E |
| px78 | CGGTCCCAGACGCCG T | Figure S1E |
| px79 | CGCTGGGGTCCATCT TTG AT | Figure S1E |

| | | |
|---|---|---|
| px80 | TGGGTCTCCGGTCCC AGT A | Figure S1E |

**Contact for Reagents and Resource sharing**

**Experimental Models and subject details**

**Human Tissue**
The Pennsylvania State University College of Medicine institutional review board approved this study (IRB Protocol number HY98-057EP-A). Methods regarding tissue procurement, mRNA isolation and processing were previously described(Schieffer et al., 2017).

**Mice**
A breeding pair of global NCLX$^{-/-}$ and wildtype C57BL/6 mice were obtained from the Jackson laboratory. NOD/SCID (NOD.CB17-*Prkdc*$^{scid}$/J) were also obtained from the Jackson laboratory. All mice were maintained in 12:12 hr light-dark cycle (22°C to 25°C) in a pathogen-free barrier facility at The Pennsylvania State University animal facility. The mice were sacrificed (age at the time of sacrifice is mentioned in the figure legends) at the end of the experiment, and organs were collected for histopathology, protein, and mRNA analysis. All experiments with mice were approved and conducted in accordance with The Pennsylvania State Institutional Animal Care and Research Advisory Committee.

**Cell culture and drug treatment**
To investigate the role of NCLX in human CRC cells, we chose two widely used CRC cell lines HCT116 and DLD1 derived from the colon of male CRC patients with colorectal carcinoma and colorectal adenocarcinoma, respectively (Ahmed et al., 2013). HCT116 and HT29 cells were cultured in McCoy's 5A media supplemented with 10% FBS and 1X antibiotic-antimycotic agents. DLD1 cells were cultured in RPMI-1640 media supplemented with 10% FBS and 1X antibiotic-antimycotic agents. All cells were kept at 37°C in a 5% $CO_2$ incubator. The cells were transfected with siRNA or plasmids using the Lipofectamine 2000 reagent, according to manufacturer protocol (Invitrogen). In experiments with 2-DG and 5-FU, cells were cultured in complete growth media McCoy's or RPMI media with 10% FBS and 1X antibiotic-antimycotic agents and treated with different concentrations of 2-DG or 5-FU for the specified time.

**Methods Details**

**Plasmids, siRNAs, shRNAs, CRISPR/Cas9, and Lentiviral infection**
The cells were seeded at 80-90% confluency at the time of transfection. The plasmids or siRNAs and Lipofectamine 2000 were diluted in Opti-MEM and mixed. The mixture was incubated for 5-10 min at room temperature and then added to the cells cultured in media with 10% serum without antibiotic-antimycotic agents. The medium of transfected cells was changed to normal media with 10% FBS and 1X antibiotic-antimycotic agents after 4-6 hrs. The expression of plasmid and siRNA was confirmed after 24 hrs and 72 hrs of transfection, respectively. To generate stable NCLX knockdown cells, shScramble, shNCLX #2, and shNCLX #3 cloned in pLKO. The lentiviral

construct was packaged into HEK293FT using the ViraPower kit (Invitrogen) and Lipofectamine 2000 (Thermofisher Scientific). HCT116 cells were infected with respective lentivirus for 72 hrs and selected with puromycin (1.5 μg/ml) for 72 hrs. The knockdown of mRNA was confirmed through RT-qPCR. Stable knockdown cells were used within two weeks, then discarded. We generated several clones of NCLX knockout (NCLX KO) in both HCT116 and DLD1 cells using the CRISPR/Cas9 system. To alleviate potential off-target effects of the CRISPR/Cas9 system, we generated several independent clones obtained with three independent guide RNAs (gRNAs; see methods) (**Figure S3A-H**). Genome sequencing and PCR on genomic DNA confirmed NCLX KO. For the case of HCT116 cells, which we generated first, NCLX KO #33 was generated using a guide RNA ($g_1$) which resulted in a single cut at nucleotide 150 in exon 1 causing a frameshift mutation and introduction of a stop codon at position 180 in the NCLX open reading frame (**Figure S3A**). NCLX KO #37 and #59 were generated using two guide RNAs ($g_1$ and $g_2$), which resulted in a double-strand cut and introduction of a stop codon in the NCLX open reading frame (**Figure S3B, C; Star method table**). For the NCLX KO clones of DLD1 generated later, we employed an advanced strategy with each knockout condition using simultaneously two guide RNAs flanking a region starting from exon 3 to the end of the NCLX gene (**Figure S3E**) to completely excise most of NCLX open reading frame. PCR on genomic DNA (**Figure S3G**) confirmed NCLX knockout in DLD1 clones. We used RT-qPCR to document the absence of mRNA in NCLX KO clones of HCT116 (**Figure S3D**) and DLD1 cells (**Figure S3H**). The positions of the primers used for qPCR are shown in **Figure S3A** and **Figure S3E** for HCT116 and DLD1 cells, respectively, and labeled as "qNCLX".

**Transcriptome sequencing (RNA-seq) and analysis**

The mRNA was isolated (minimum 3 μg) using the RNeasy Mini Kit. A fragmentation buffer (Ambion) was used to short fragment the mRNA. These short-fragmented mRNA were used as a template to synthesize double-stranded cDNA. The cDNA was end-repaired and ligated to Illumina adapters. Further, to generate the library, these cDNAs were size selected (~250 bp) on an agarose gel, and PCR amplified. To sequence the cDNA, the prepared cDNA library was fed to the Illumina HiSeq 2500 sequencing platform (Berry Genomics). The expression level of each transcript of a gene was quantified using read counts with HTseq. After removing gene identifiers with zero read counts in each sample, the biomaRt R package (Durinck et al., 2009) was used to convert Ensemble gene identifiers to gene symbols using the January 2019 archive (http://jan2019.archive.ensembl.org). Identifiers corresponding to multiple gene symbols were removed. The edgeR R package (McCarthy et al., 2012; Robinson et al., 2010) was used to identify lowly expressed genes based on thresholds defined using counts per million reads mapped. This yielded expression data for a total of 10,179 genes.

Exploratory analyses were performed after utilizing the VST (variance stabilizing transformation) function in the DESeq2 R package (Love et al., 2014) to apply a variance stabilizing transformation to the read count data. The results strongly suggested the presence of a batch effect. After applying the voom transformation (Law et al., 2014) to the read counts, the limma R package (Ritchie et al., 2015) was used to identify differentially expressed genes based on a q-value threshold of 0.05. The limma design matrix included batch information as a factor.

**Gene set enrichment analysis (GSEA)**

The gene set enrichment analysis (GSEA) software (Subramanian et al., 2005) was used to perform pathway analyses based on the limma output. Briefly, genes in the limma output were ordered according to the $t$ test statistic. Then a GSEA pre-ranked analysis was performed using the Hallmark Gene Sets in the Molecular Signatures Database (http://software.broadinstitute.org/gsea/msigdb/index.jsp).

**TCGA analysis**

RNA-sequencing (RNA-seq) based gene expression data for both tumor and normal samples, as well as clinical data for the combined TCGA colon adenocarcinoma and rectal adenocarcinoma (COADREAD) cohort were downloaded from the Broad Institute's Firehose GDAC (https://gdac.broadinstitute.org/). Gene expression was quantified as log2(normalized RSEM + 1). Somatic gene mutation data was obtained from (Ellrott et al., 2018) after restricting to samples in the RNA-seq cohort. Wilcoxon rank-sum tests and Kruskal-Wallis tests were used to compare expression values of NCLX/SLC24A6 across groups defined by the clinical and mutation data.

**Reverse transcription-quantitative polymerase chain reaction (RT-qPCR)**

Human colorectal cancer tissues utilized for mRNA extraction were obtained from the Penn State Hershey medical center with IRB approval. Briefly, mRNA were isolated from either tissues or cells. The cells were washed with cold PBS, and RNA was isolated using the RNeasy Mini Kit. The quality and quantity of RNA were analyzed using NanoDrop2000 (Thermo Scientific, Wilmington, DE, USA). High-Capacity cDNA Reverse Transcription Kit (Applied Biosystems, Foster City, CA) was used to make cDNA. The cDNA was further used for quantitative real-time PCR using the SYBR select master mix (Thermo Scientific) according to the manufacturer protocol. The reactions were run in technical duplicates using the Quant Studio 3 system (Applied Biosystems). The expression levels of genes were normalized to at least two housekeeping gens, GAPDH or Tubulin or NONO (specified in the figure legends) using the $2^{-\Delta Ct}$ method.

**Western blot**

The cells were cultured with 80-90% confluency and were washed with chilled PBS and lysed in 100 µl RIPA buffer (Sigma) containing Halt protease and phosphatase Inhibitor. 30-50 µg of the protein was loaded on a 4-12% gel (NuPAGE Bis-Tris precast gels, Life Technologies) and transferred to a polyvinylidene difluoride membrane. The membrane was incubated in Odyssey blocking buffer for 1 hr at room temperature for blocking and was incubated overnight at 4°C in primary antibody diluted in Odyssey blocking buffer. The primary antibodies were used are mentioned in the key resource table, and their dilutions are as follows: anti-HIF1α (1:500), all other antibodies were used at 1:1000 dilution. The next day, the membrane was washed with 0.1% TBS-T for 5 min (three times) at room temperature, followed by incubation in IRDye for 2 hrs at room temperature. The IRDyes used are mentioned in the key resource table, and they were diluted in Odyssey blocking buffer (IRDye 680RD Goat anti-Mouse at 1:10,000 dilution and IRDye 800CW Donkey anti-Rabbit at 1:5000). The membrane was washed with 0.1% TBS-T for 5 minutes (three times) at room temperature. Then the blot was imaged in Odyssey CLx Imaging System (LI-COR, NE, USA). Densiometric analysis of the bands on the membrane was performed using ImageJ software.

**Immunofluorescence microscopy**

The cells were cultured in 6 well plates on glass coverslip at 60-70% confluency. The next day, the cells were washed with cold PBS and fixed with chilled 100% Methanol for 30 min. The cells

were washed with chilled PBS (3X for 5 min). The cells were incubated in an anti-cleaved caspase-3 antibody (1:250). Secondary Alexa Fluor 594 (1:5000) was used to stain the cells.
For live-cell imaging and mitochondrial staining, the cells were stained with MitoTracker Green FM (50 nM in complete media for 30 min at 37°C), TMRM (100 nM in complete media for 30 min at 37°C) or MitoSOX Red (2.5 μM in HBSS for 30 min at 37°C) and Hoechst (1 μM in HBSS for 5 min). The images were acquired using the Leica SP8 confocal microscope. The fluorescence intensity was quantified using LAS X (Leica microsystems) software.

**Mitochondrial $Ca^{2+}$ measurements**
To measure mitochondrial $Ca^{2+}$ using Rhod-2 AM (543 nm/580-650 nm) dye, the cells were cultured at 50-60% confluency. The cells were washed with media without FBS and antibiotic-antimycotic agents. Then the cells were incubated in media containing 1 μM Rhod-2 AM (without FBS and antibiotic-antimycotic agents) at 37°C for 30 min. The cells were washed with media and were loaded with MitoTracker Green FM (50 nM in media for 30 min at 37°C) for photobleaching and focus corrections. After loading with MitoTracker Green, cells were washed and kept in HBSS (HEPES-buffered saline solution (140 mM NaCl, 1.13 mM $MgCl_2$, 4.7 mM KCl, 2 mM $CaCl_2$, 10 mM D-glucose, and 10 mM HEPES, adjusted to pH 7.4 with NaOH)) containing 5 mM $CaCl_2$ for imaging. The cells were stimulated with 300 μM ATP in HBSS containing 5 mM $CaCl_2$. The time-lapse images were acquired using the Leica TCS SP8 confocal microscope equipped with a 63X oil objective. The images were acquired every 5-sec intervals for 10-15 min. The fluorescence intensity was quantified using LAS X (Leica microsystems) software.

**Intracellular $Ca^{2+}$ measurements**
Intracellular $Ca^{2+}$ imaging was performed described previously (Emrich et al., 2019; Zhang et al., 2019). Briefly, the cells were cultured on glass coverslips with 50-60% confluency. Next day the cells were washed with fresh media, and the glass coverslip with cells were mounted on a Teflon chamber. Further, the cells were incubated at 37°C for 45 min in culture media containing 4 μM Fura-2/acetoxymethyl ester (Molecular Probes, Eugene, OR, USA). Then the cells were washed and kept in HEPES-buffered saline solution (140 mM NaCl, 1.13 mM $MgCl_2$, 4.7 mM KCl, 2 mM $CaCl_2$, 10 mM D-glucose, and 10 mM HEPES, adjusted to pH 7.4 with NaOH) for at least 10 min before $Ca^{2+}$ measurements were made. Time-lapse imaging was performed, and the change in fluorescence of Fura-2 was recorded and analyzed with a digital fluorescence imaging system (InCyt Im2; Intracellular Imaging Inc., Cincinnati, OH, USA). An ROI was drawn around each cell, and 340/380 ratio of each cell was calculated.

**Transmission Electron Microscopy (TEM)**
The HCT116, DLD1, NCLX KO HCT116, and NCLX KO DLD1 cells were cultured at 80-90% confluency and fixed with 1% glutaraldehyde in 0.1 M sodium phosphate buffer, pH 7.3. After fixation, the cells were washed with 100 mM Tris (pH 7.2) and 160 mM sucrose for 30 min. The cells were washed again twice with phosphate buffer (150 mM NaCl, 5 mM KCl, 10 mM $Na_3PO_4$, pH 7.3) for 30 min, followed by treatment with 1% $OsO_4$ in 140 mM $Na_3PO_4$ (pH 7.3) for 1 hr. The cells were washed twice with water and stained with saturated uranyl acetate for 1 h, dehydrated in ethanol, and embedded in Epon (Electron Microscopy Sciences, Hatfield, PA). Roughly 60 nm section were cut and stained with uranyl acetate and lead nitrate. Further, the stained grids were analyzed using a Philips CM-12 electron microscope (FEI; Eindhoven, The

Netherlands) and photographed with a Gatan Erlangshen ES1000W digital camera (Model 785, 4 k 3 2.7 k; Gatan, Pleasanton, CA). Morphometric analysis of mitochondrial and mitophagy was analyzed by double-blinded independent observers in at least 10 different micrographs per condition. The mitochondria without intact inner mitochondrial membrane and cristae were classified as damaged, and their number was counted in each cell and divided by the total number of mitochondria in that cell to calculate % damaged mitochondria. The mitochondrial area was measured by manually tracing the outer mitochondrial membrane and using the measure function of NIH ImageJ software. Cristae per mitochondria were calculated by counting the number of intact cristae in each mitochondrion.

**Mitochondrial membrane potential and Mitochondrial ROS measurements**

Cells were cultured in 6-well plates at 50-60% confluency. The next day, 1 x $10^6$ cells were harvested and loaded with Tetramethyl rhodamine (TMRM) dye. The cells were stained with 100 nM TMRM dye in complete growth media and kept at 37°C in 5% $CO_2$ for 20-30 min. CCCP (50 µM) was used as a positive control. To measure mitochondrial ROS, 1 x $10^6$ cells were stained with MitoSOX (2.5 µM in HBSS at 37°C for 30 min). Antimycin (50 µM) was used as a positive control. The intensity of staining was measured on an LSRII flow cytometer using FACSDiva software (BD Biosciences) and analyzed with FlowJo software (Tree Star).

**Cell proliferation assays**

The cells were harvested, and 3000-5000 cells were plated in each well of 96 well plates. The cells were kept at 37°C in 5%$CO_2$ for 4 hrs to allow the cells to adhere to the plate. The CyQUANT-NF dye was diluted in HBSS buffer, and 100µl of the mixture was added in each well. The plate was kept at 37°C in 5% $CO_2$ for 1 hr. The fluorescence intensity (~485 nm/~530 nm) was measured using FlexStation 3 Multimode Plate Reader (VWR). To analyze the effect of 2DG and 5-FU on proliferation, different doses of the drugs were added in the culture media and cells were grown for the indicated amount of time. The fluorescence intensity of the dye was recorded after the drug treatment. The normalized intensity was calculated using the following formula:

Normalized intensity= $(I_t-I_b)/(I_0-I_b)$

$I_t$= intensity at the given time

$I_b$= background intensity

$I_0$= intensity at time zero

**Zymography**

Briefly, the cells were harvested and cultured in 6-well plates. Once the culture becomes 80-90% confluent, 20 µl of the media was taken and mixed with Tris-Glycine-SDS sample buffer and loaded in a Novex Zymogram gel and Tris-glycine SDS running buffer was used to run the gel. After electrophoresis, the gel was kept in Novex Zymogram renaturing buffer for 30 min and transferred to Novex Zymogram Developing buffer and kept at 37°C overnight. The next day the gel was stained with SimplyBlue™ Safe Stain and imaged using FluorChem M imager. The images were analyzed using ImageJ.

***In vitro* migration and invasion assays**

Migration was measured using two different methods a transwell migration assay and wound healing assay. For the transwell migration assay using FluoroBlok, the cells were serum-starved for 24 hrs, and 1,000 cells were plated on the chamber in 200 µl media without FBS. The cells were stimulated to migrate towards preconditioned media with 10% FBS. After 8 hrs of migration, the upper part of the membrane was cleaned, and the lower part was stained with DAPI, and the intensity of DAPI was measured using FlexStation 3 Multimode Plate Reader (VWR).

For the wound healing (gap closure) assay, cells were serum-starved for 24 hrs to synchronize the cell cycle. Then 50,000 cells were plated in ibidi-silicone insert with a defined cell-free gap. After 24 hrs, the inserts were lifted, and the gap between two migrating fronts was measured at 0 hrs, 12 hrs, and 24 hrs using Zeiss inverted fluorescence microscope equipped with 5X air objective. The area between the two migrating fronts was measured using ImageJ (NIH, Bethesda). To analyze the effect of 2-DG and 5-FU on migration, a specific dose of the drugs was added in the culture media at the time of removal of the silicon insert, and cells were grown for the indicated amounts of time. Fresh media with the drug was added every 12 hr. The following formula was used to calculate % gap closure-

% Gap closure= $(A_0-A_t/A_0)/100$

$A_0$= area of gap measured immediately after lifting the insert

$A_t$= area of gap measured after the indicated time of migration

Cell invasion was measured using the BioCoat™ Tumor Invasion Plate with 8 µm pore size inserts, which were coated with growth factor-reduced Matrigel. The cells were serum-starved for 24 hrs, and a total of 5,000 cells were plated on the chamber in 200 µl media without FBS. The cells were stimulated to invade towards preconditioned media with 10% FBS. After 24 hrs invasion, the upper part of the membrane was cleaned, and the lower part was stained with DAPI, and the intensity of DAPI was measured using FlexStation 3 Multimode Plate Reader (VWR). The percentage of invasion was calculated using the following formula-

% invasion= (Fluorescence of invading cells - Fluorescence of blank chamber)/ Fluorescence of migrating cells - Fluorescence of blank chamber) X100

**Extracellular acidification rate (ECAR) and Oxygen consumption rate (OCR)**

The ECAR, OCR, and % metabolite dependency was measured using the Seahorse XFp Extracellular Flux Analyzer (Seahorse Bioscience). All experiments were performed according to the manufacturer's protocol. Seahorse XFp Glycolysis Stress Test Kit, Seahorse XFp Cell Mito Stress Test Kit, and Seahorse XFp Mito Fuel Flex Test Kit (Agilent Technologies) were used to measure ECAR, OCR, and % dependency respectively. Briefly, cells were harvested, and 30,000 cells were plated per well and cultured in a Seahorse XFp cell culture microplate with complete growth media for 10-12 hr. The next day the cells were washed with seahorse XF DMEM media (pH 7.4). The plate with cultured cells was placed in the Seahorse XFp Extracellular Flux Analyzer, and baseline measurements were recorded. For ECAR measurements, oligomycin, and 2-DG were sequentially injected into each well at the indicated time points. Similarly, for OCR measurements, oligomycin, FCCP (p-trifluoromethoxy carbonyl cyanide phenylhydrazone), and antimycin A (Rote/AA) were sequentially injected. For % dependency measurements, UK5099, BPTES, and Etomoxir were sequentially injected. The results obtained for ECAR, OCR, and Mito Fuel Flex Test were normalized to cell number. Data were analyzed using Seahorse XFp Wave software. The results for OCR are reported in pmols/minute, ECAR in mpH/minute, and Mito Fuel Flex Test in % dependency.

**Glucose and Lactate measurements**

To measure glucose and lactate in media, 100,000 cells were plated in each well of 6-well plates with 2 ml of complete growth media. After 24 hrs, 400 µl of the media were taken from each well

of the 6-well plates and divided into 4 wells in a 96-well plate (quadruplicate for each NCLX KO clone and respective control). Then the plate was kept in YSI 7100 multichannel biochemistry analyzer (YSI Life Sciences) to measure glucose and lactate levels in the media. Fresh media was used to measure the basal level of glucose and lactate. After measurements were completed, cells were harvested, and the protein content was measured using the BCA assay. This experiment was performed at least three times independently. The following formulas were used to calculate glucose consumption and lactate generation:
Glucose consumption= (Glucose in fresh media - Glucose in the media with cells)/ protein content
Lactate generation= (Lactate in the media with cells - Lactate in fresh media)/ protein content

**Mouse xenograft experiments**
Mice were anesthetized using isoflurane, and after proper sterile preparation of the abdomen, a small (4-8 mm) incision was made by sharp sterile blade over the left upper quadrant of the abdomen. The peritoneal cavity was carefully exposed, and the spleen was located. Further, the spleen was carefully exteriorized on the sterile field around the incision area, and 500,000 cells suspended in 50 ml McCoy's (10 or 20% FBS) were then injected into the spleen via a 1 ml insulin syringe. A small bleb was observed while injecting the cells in the spleen, confirming that the cells were successfully injected. The spleen was carefully placed back into the peritoneal cavity. Both the peritoneal cavity and skin were closed with sterile absorbable suture. The mice injected with luciferin (100 µg/ml) and imaged in the IVIS system every week till the time of sacrifice.

**Colitis-associated colon cancer model and histology**
6-8-week-old mice were given a single Azoxymethane (AOM) injection via IP. One week after AOM injection, the mice were given three cycles of dextran sodium sulfate (DSS), which was supplemented in drinking water (1.5 % m/v, MW 36-50kDa, MP Biochemicals) *ad libitum* for five days. The DSS cycles were intermixed with two weeks of normal autoclaved drinking water. Mice were sacrificed, and the colons were collected ten weeks after the last DSS cycle. The collected colons were flushed with PBS to clear feces and photographed. Before dissection, tumors were scored according to their number and size. PBS-rinsed colons were either snap-frozen for further analysis or fixed in 10% neutral buffered formalin and sectioned for histological analysis.

**Quantification and statistical analysis**
Data are represented as mean ±SEM and analyzed using Origin pro 2019b (Origin lab). In the box plot, the box represents the 25$^{th}$ to 75$^{th}$ interquartile range, midline in the box represents the median, and the solid square box represents mean data points. To test single variables between two groups, paired t-test or Kruskal-Wallis ANOVA was performed (specified in each figure legend). One-way ANOVA followed by post hoc Tukey's test was used for comparison between multiple conditions and the control group. The p-value < 0.05 was considered to be significant and is presented as *p < 0.05, **p < 0.01, or ***p < 0.001.

**Legends to supplementary figures**

**Figure S1: Loss of NCLX reduces tumor number and size in the colitis-associated cancer model**

(**A**) TCGA data analysis showing NCLX mRNA levels in tumors of male and female COADREAD patients.

**(B, C)** TCGA data analysis showing a comparison of NCLX mRNA levels in COAD patients based on age **(B)** and the origin of adenocarcinoma **(C)**.

**(D)** Cartoon depicting the position of the nucleotides deleted from the NCLX gene in NCLX $^{-/-}$ mice. The red color depicts the deleted region, the light green bar represents the NCLX gene, and the dark green boxes represent the coding regions in the exons.

(**E**) Cartoon representing the annealing position of the primer pair sets used for genotyping NCLX$^{-/-}$ mice, and products of the PCR performed on genomic DNA from wild type (WT), heterozygotes (NCLX$^{-/+}$), and NCLX$^{-/-}$ mice tails.

(**F, G**) Magnified image of a colon from an NCLX$^{-/-}$ mouse and a littermate control mouse where the tumors are marked by red arrows **(F)** and colons isolated at day 78 from fifteen NCLX$^{-/-}$ mice and control littermate mice subjected to AOM/DSS treatment **(G)**.

In TCGA analysis, each data point represents individual patients. Kruskal-Wallis ANOVA was used to calculate statistics. *$p<0.05$, **$p<0.01$, and ***$p<0.001$

**Figure S2: Xenografts of HCT116 cells and HCT116 NCLX KO #33 cells in NOD/SCID**

Bioluminescence images of the remaining male NOD/SCID mice injected with luciferase-tagged HCT116 cells or HCT116 NCLX KO #33 cells.

**Figure S3: Deletion of NCLX results in reduced proliferation and increased migration and invasion**

(**A-D**) CRISPR/Cas9 knockout strategy of the NCLX gene in NCLX KO clones #33 (**A**), #37 (**B**), and #59 (**C**) with g1 and g2 representing the cut site and red box non-translated region. Location of primers used for RT-qPCR are shown in (**A**), and NCLX mRNA levels in clones of HCT116 NCLX KO cells compared to control HCT116 cells **(D)**.

(**E-H**) The binding site of guide RNA (gRNA) g3 and g4 on the NCLX gene, the subsequent deletion, and location of primers used for RT-qPCR (**E**). The position on the NCLX gene of the genotyping primers used to screen gene knockout of NCLX KO clones **(F)**, PCR data on genomic DNA showing amplified products with primer pair sets **(G),** and RT-qPCR data from DLD1 cells and clones of DLD1 NCLX KO cells normalized to tubulin **(H)**.

(**I**) RT-PCR data showing NCLX mRNA relative to tubulin in HCT116 cells transfected with shRNA against NCLX (shNCLX #2, shNCLX #3, and shNCLX #2+3) and normalized to scramble control.

(**J**) Comparison of proliferation between HCT116 cells infected with scramble shRNA and two different NCLX shRNA sequences.

(**K, L**) Representative image of HCT116 cells and HCT116 NCLX KO #33 cells stained with Hoechst and anti-cleaved caspase-3 antibody **(K)**, and quantification of the percentage of cells showing cleaved caspase-3 staining **(L)**. Each data point represents one replicate with each replicate representing the average values from at least three image fields. Total cell counted, HCT116 cells (n = 100), and HCT116 NCLX KO #33 cells (n = 135).

(**M, N**) Quantification of % gap closure for HCT116 cells and clones of HCT116 NCLX KO cells after 6 hr **(M)**, and with and without mitomycin (300 μM) treatment for 12 and 24 hr **(N)**.
(**O**) Quantification of *in vitro* invasion of DLD1 cells and clones of DLD1 NCLX KO cells.
(**P, Q**) Western blots probed with anti-MMP1, anti-MMP9, and anti-GAPDH antibodies **(P)**. Quantification of protein band intensities between DLD1 cells and DLD1NCLX KO clones relative to GAPDH (**Q**).
(**R, S**) Western blots probed with anti-MMP1, anti-MMP9, and anti-GAPDH antibodies (**R**). Quantification of protein band intensities between HCT116 cells infected with scramble shRNA and NCLX shRNA #2, #3, and #2+3 relative to GAPDH (**S**).
All experiments were replicated ≥three times with similar results. Statistical significance was calculated using one-way ANOVA followed by a post-hock Tukey test, except for L, O, Q, and S, where paired t-test was used. *p<0.05, **p<0.01, and ***p<0.001

**Figure S4: Loss of NCLX causes reduced mtCa$^{2+}$ extrusion**
(**A-D**) ATP-stimulated mitochondrial Ca$^{2+}$ uptake and Ca$^{2+}$ extrusion measured with the dye Rhod-2 in HCT116 (n=72), HCT116 NCLX KO #37 (n=118) cells (**A**), HCT116 (n=198), HCT116 NCLX KO #59 (n=256) cells (**B**). Similar recordings in DLD1 (109), DLD1 NCLX KO #24 (n=90) cells (**C**), DLD1 (n=179) and DLD1 NCLX KO #32 (n=107) cells (**D**). The Rhod-2 intensity was normalized to mitoGFP intensity. The traces are represented as mean±S.E.
(**E**) RT-qPCR data depicting NCLX mRNA levels in HCT116, DLD1, and HT29 cells transfected with siRNA against NCLX compared to their respective siRNA control-transfected cells.
(**F-N**) ATP-stimulated mitochondrial Ca$^{2+}$ uptake and Ca$^{2+}$ extrusion measured with the dye Rhod-2 in scramble HCT116 (n=48) and siNCLX HCT116 (n=38) (**F**), its Ca$^{2+}$ uptake **(G)**, Ca$^{2+}$ extrusion **(H)**, scramble DLD1(n=16) and siNCLX DLD1(n=17) **(I)**, its Ca$^{2+}$ uptake **(J)**, Ca$^{2+}$ extrusion **(K),** scramble HT29 (n=14) and siNCLX HT29 (n=8) (**L**) and its Ca$^{2+}$ uptake **(M)**, Ca$^{2+}$ extrusion **(N).** The traces are represented as mean±S.E.
(**O, P**) Representative trace showing cytosolic Ca$^{2+}$ in HCT116 (n=110) and HCT116 NCLX KO #33 (100) cells measured by Fura-2 in response 300 μM ATP in nominally Ca$^{2+}$-free external solution and subsequent addition of 10 mM Ca$^{2+}$. The traces are represented as mean±S.E **(O)**, and peak SOCE after addition of 10 mM Ca$^{2+}$ **(P)**.
All experiments were replicated ≥three times with similar results. Kruskal-Wallis ANOVA was used to calculate statistical significance. *p<0.05, **p<0.01, and ***p<0.001

**Figure S5: Loss of NCLX causes mitochondrial damage**
(**A-C**) Representative transmission electron micrographs of mitochondria in HCT116, HCT116 NCLX KO #33 cells (**A**), HCT116 NCLX KO #37 cells, HCT116 NCLX KO #59 cells (**B**), DLD1 cells, DLD1 NCLX KO#06 cells (**C**). The red arrows point to damaged mitochondria. The magnified image shows mitochondria encapsulated in a phagosome **(A)**.
(**D, E**) Western blots showing levels of LC3B and p62 in control DLD1 cells NCLX KO clones of DLD1 cells **(D)**, and quantification of band intensity relative to loading control GAPDH **(E)**.
(**F, G**) Western blots probed with anti-LC3B and anti-p62 in HCT116 cells infected with control scrambled shRNA or three different NCLX shRNA conditions (**F**). Quantification of band intensity relative to HSC70 (**G**).
(**H-K**) Measurement of OCR in DLD1 cells and DLD1 NCLX KO clone #06 (**H, I**) and from DLD1 NCLX KO clone #24, and clone #32 and (**J, K**). Quantification of Basal respiration (Basal

Res), Spare respiratory capacity (Spare Res Cap), ATP generation (ATP Gen), and maximal respiration (Max Res) (**I, K**).

**(L, M)** Western blots probed with a cocktail of antibody against electron transport chain complexes, including ATP5A for complex V, UQCRC2 for complex III, MTCO1 for complex IV, SDHB for complex II, and NDUFB8 for complex I **(L)**, and quantification of their band intensity **(M)**.

(**N, O**) Representative images of mitoSox, and Hoechst fluorescence in siRNA control-transfected HCT116, HT29, and DLD1 cells with their respective siRNA NCLX-transfected cells **(N)**. The mitoSox fluorescence intensity was quantified using Flow cytometry (**O**). Scale bar, 10 µm

All experiments were performed ≥three times. The statistics were calculated by paired t-test unless mentioned otherwise, *p≥0.05, **p≥0.01, ***p≥0.001

**Figure S6: Abrogation of NCLX function leads to transcriptional reprogramming.**

(**A**) Heat map of expression of glycolytic genes that are significantly different between HCT116 cells and HCT116 NCLX KO #33 cells.

(**B, C**) Gene set enrichment analysis (GSEA), and of HCT116 cells, and HCT116 NCLX KO #33 shows enrichment of positive correlation of hallmark of apoptotic genes **(B)** and heat map of differentially expressed apoptosis-related genes in NCLX KO #33 cells **(C)**.

(**D**) Heat map of differentially regulated cell cycle genes in HCT116 cells and HCT116 NCLX KO #33 cells.

(**E**, **F**) Gene set enrichment analysis (GSEA) (**E**) and heat map (**F**) of significantly different genes between HCT116 cells and HCT116 NCLX KO #33 cells.

**(G)** The proliferation of HCT116 cells and HCT116 NCLX KO #33 cells in the presence of 0 µM, 1 µM, 10 µM, and 25 µM 5-FU. n=3

Statistical significance was calculated one-way ANOVA followed by a post-hock Tukey. *p<0.05, **p<0.01, and ***p<0.001.

**Figure S7: NCLX KO CRC cells show chemoresistance to 5-FU**

(**A-H**) RT-qPCR data plotted as the $2^{-\Delta Ct}$ mRNA value of SLC7A11 (**A**), GCLM (**B**) in clones of HCT116 NCLX KO cells normalized to control HCT116 cells, and SLC7A11 (**C**), GCLM (**D**) NANOG (**E**), Sox2 (**F**), Fox3 (**G**), and Oct4 (**H**) in clones of DLD1 NCLX KO cells relative to tubulin and normalized to control DLD1 cells.

(**I**) Schematic representation of HIF1α induced transcriptional activation of SLC7A11 and GCLM, leading to increased transcription of stem cell genes NANOG, Oct4, and Sox2 in CRC cells.

**(J)** Measurement of IC50 of 5-FU for HCT116 and HCT116 NCLX KO #33 cells after treating the cells with 0 µM, 1 µM, 10 µM, and 25 µM 5-FU for 72 hrs.

(**K**) The proliferation of DLD1 cells and clones of DLD1 NCLX KO cells in the absence and presence of 10 µM 5-FU.

(**L, M**) Quantification of migration of HCT116, clones of HCT116 NCLX KO cells **(L)**, and DLD1 cells and clones of DLD1 NCLX KO cells **(M)** in the presence and absence of 10 µM 5-FU.

(**N-Q**) Immunoblot for phospho-AMPK, total AMPK **(N)**, phosphor-S6K, total S6K **(P)** and loading control GAPDH protein levels, and quantification of band intensity normalized to GAPDH from HCT116 cells and HCT116 NCLX KO #33, #37, and #59 clones (**O, Q**).

All the experiments were replicated ≥ three times with similar results. Statistical significance was calculated using the paired t-test, except for K, L, and M, where one-way ANOVA followed by a post-hock Tukey test was performed. *p<0.05, **p<0.01, and ***p<0.001.

**Figure S8: Loss of NCLX enhances glycolysis and diminishes mitochondrial respiration**

(**A-D**) Western blots for HK2, ALDOA, LDHA, with GAPDH as a loading control from HCT116, HCT116 NCLX KO #37, #59 **(A)**, DLD1, clones of DLD1 NCLX KO cells **(C)**, and quantification of band intensity normalized to GAPDH **(B, D)**.

(**E, F**) Western blots probed with anti-HK2, anti-ALDOA, anti-LDHA, and anti-HSC70 loading control (**E**), and quantification of band intensity **(F)** from HCT116 cells transfected with scramble and two different constructs of shNCLX.

(**G**) RT-qPCR data plotted as $2^{-\Delta Ct}$ mRNA values relative to tubulin and normalized to control, for Glucose-6-phosphate dehydrogenase (G6PD), Phosphogluconate dehydrogenase (PGD), and Transketolase (TKT) in HCT116 cells and HCT116 NCLX KO #33 cells.

(**H**) ECAR measurement from DLD1 cells and clones of DLD1 NCLX KO cells.

(**I**) The proliferation of DLD1 cells and clones of DLD1 NCLX KO cells in the presence and absence of 2.5 mM 2-DG.

(**J**) RT-qPCR analysis of NCLX mRNA in CRC patient tumors, adjacent normal tissue, HCT116, and DLD1 cells.

All experiments were performed ≥three times. The statistics were calculated by One-way ANOVA followed by post-hock Tukey's test except for B, D, and F where paired t-test was performed, *p≥0.05, **p≥0.01, ***p≥0.001

#### References

Ahmed, D., Eide, P. W., Eilertsen, I. A., Danielsen, S. A., Eknæs, M., Hektoen, M., Lind, G. E., and Lothe, R. A. (2013). Epigenetic and genetic features of 24 colon cancer cell lines. Oncogenesis *2*, e71-e71.
Durinck, S., Spellman, P. T., Birney, E., and Huber, W. (2009). Mapping identifiers for the integration of genomic datasets with the R/Bioconductor package biomaRt. Nat Protoc *4*, 1184-1191.
Ellrott, K., Bailey, M. H., Saksena, G., Covington, K. R., Kandoth, C., Stewart, C., Hess, J., Ma, S., Chiotti, K. E., McLellan, M.*, et al.* (2018). Scalable Open Science Approach for Mutation Calling of Tumor Exomes Using Multiple Genomic Pipelines. Cell Systems *6*, 271-281.e277.
Emrich, S. M., Yoast, R. E., Xin, P., Zhang, X., Pathak, T., Nwokonko, R., Gueguinou, M. F., Subedi, K. P., Zhou, Y., Ambudkar, I. S.*, et al.* (2019). Cross-talk between N-terminal and C-terminal domains in stromal interaction molecule 2 (STIM2) determines enhanced STIM2 sensitivity. J Biol Chem *294*, 6318-6332.
Law, C. W., Chen, Y., Shi, W., and Smyth, G. K. (2014). voom: precision weights unlock linear model analysis tools for RNA-seq read counts. Genome Biology *15*, R29.
Love, M. I., Huber, W., and Anders, S. (2014). Moderated estimation of fold change and dispersion for RNA-seq data with DESeq2. Genome Biol *15*, 550.
McCarthy, D. J., Chen, Y., and Smyth, G. K. (2012). Differential expression analysis of multifactor RNA-Seq experiments with respect to biological variation. Nucleic Acids Res *40*, 4288-4297.
Ritchie, M. E., Phipson, B., Wu, D., Hu, Y., Law, C. W., Shi, W., and Smyth, G. K. (2015). limma powers differential expression analyses for RNA-sequencing and microarray studies. Nucleic Acids Res *43*, e47.
Robinson, M. D., McCarthy, D. J., and Smyth, G. K. (2010). edgeR: a Bioconductor package for differential expression analysis of digital gene expression data. Bioinformatics *26*, 139-140.
Schieffer, K. M., Choi, C. S., Emrich, S., Harris, L., Deiling, S., Karamchandani, D. M., Salzberg, A., Kawasawa, Y. I., Yochum, G. S., and Koltun, W. A. (2017). RNA-seq implicates deregulation of the immune system in the pathogenesis of diverticulitis. Am J Physiol Gastrointest Liver Physiol *313*, G277-G284.
Subramanian, A., Tamayo, P., Mootha, V. K., Mukherjee, S., Ebert, B. L., Gillette, M. A., Paulovich, A., Pomeroy, S. L., Golub, T. R., Lander, E. S., and Mesirov, J. P. (2005). Gene set enrichment analysis: a knowledge-based approach for interpreting genome-wide expression profiles. Proc Natl Acad Sci U S A *102*, 15545-15550.
Zhang, X., Pathak, T., Yoast, R., Emrich, S., Xin, P., Nwokonko, R. M., Johnson, M., Wu, S., Delierneux, C., Gueguinou, M.*, et al.* (2019). A calcium/cAMP signaling loop at the ORAI1 mouth drives channel inactivation to shape NFAT induction. Nature Communications *10*, 1971.

**Figure S1**

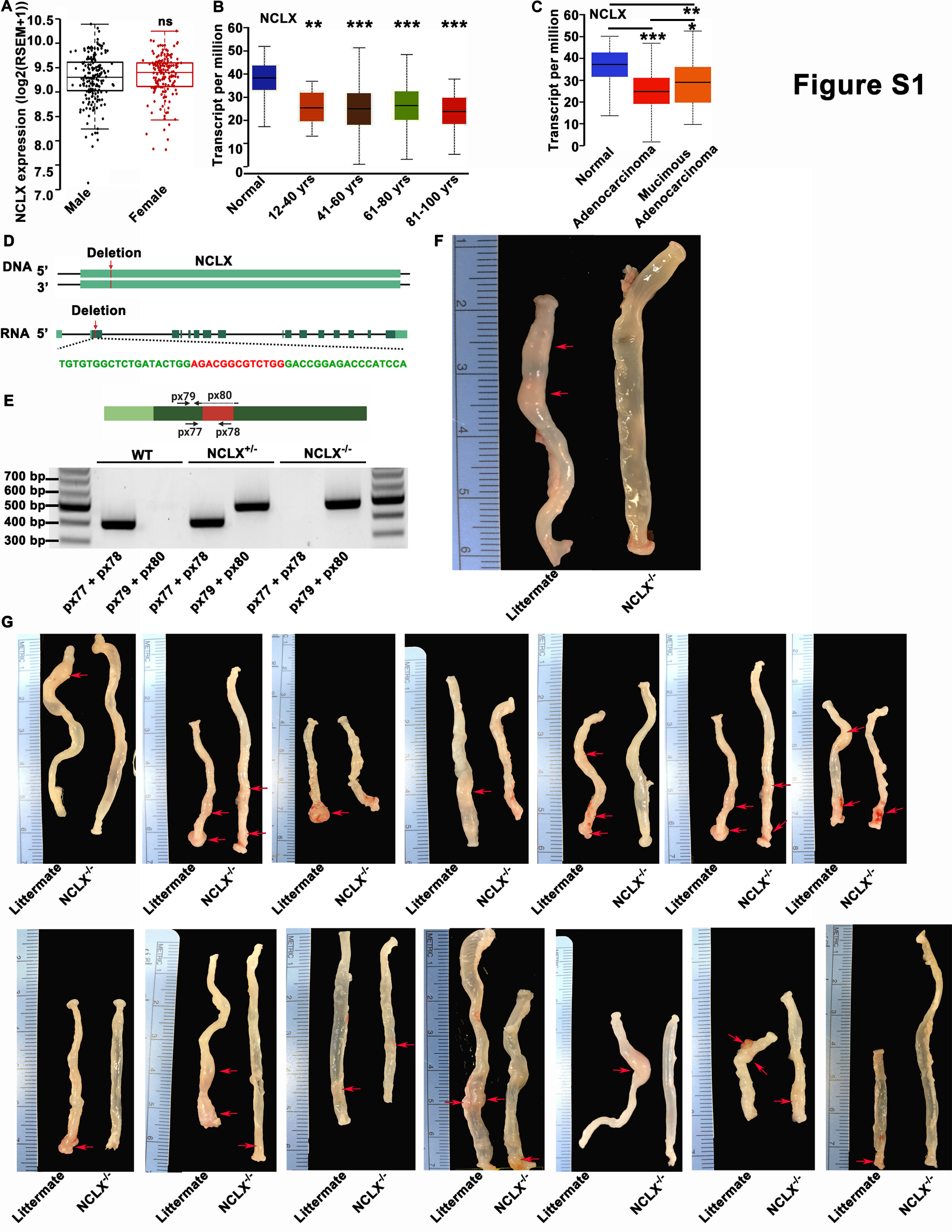
A
NCLX expression (log2(RSEM+1))
10.5
10.0
9.5
9.0
8.5
8.0
7.5
7.0
ns
Male
Female
B
NCLX
Transcript per million
60
50
40
30
20
10
0
** *** *** ***
Normal
12-40 yrs
41-60 yrs
61-80 yrs
81-100 yrs
C
NCLX
Transcript per million
60
50
40
30
20
10
0
***
**
*
Normal
Adenocarcinoma
Mucimous Adenocarcinoma
D
Deletion
NCLX
DNA 5'
3'
Deletion
RNA 5'
TGTGTGGCTCTGATACTGGAGACGGCGTCTGGGACCGGAGACCCATCCA
E
px79
px80
px77
px78
WT
NCLX+/-
NCLX-/-
700 bp
600 bp
500 bp
400 bp
300 bp
px77 + px78
px79 + px80
px77 + px78
px79 + px80
px77 + px78
px79 + px80
F
Littermate
NCLX-/-
G
Littermate
NCLX-/-

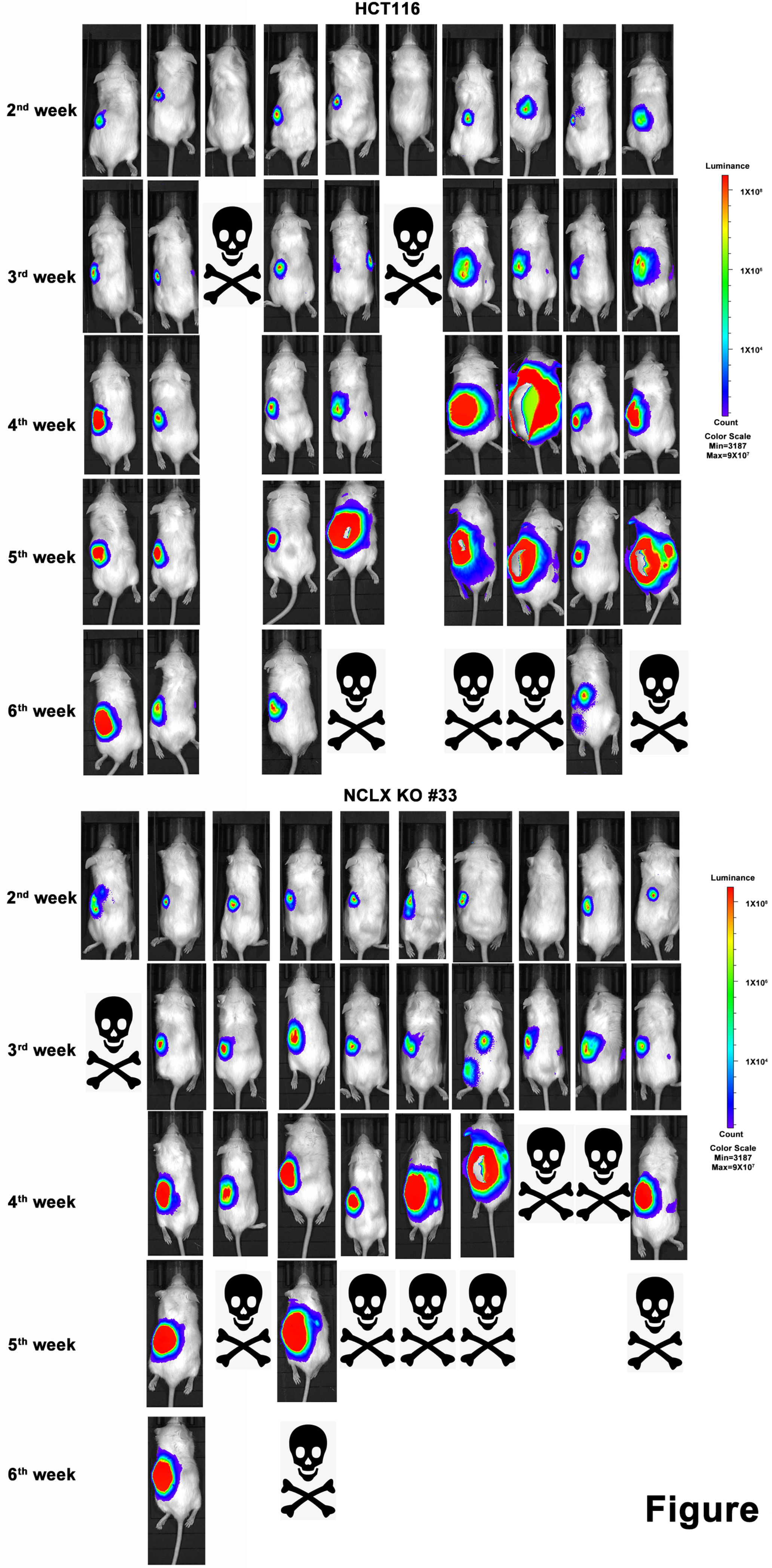
HCT116
2nd week
3rd week
4th week
5th week
6th week
Luminance
$1X10^8$
$1X10^6$
$1X10^4$
Count
Color Scale
Min=3187
Max=$9X10^7$
NCLX KO #33
2nd week
3rd week
4th week
5th week
6th week
Luminance
$1X10^8$
$1X10^6$
$1X10^4$
Count
Color Scale
Min=3187
Max=$9X10^7$

**Figure S2**

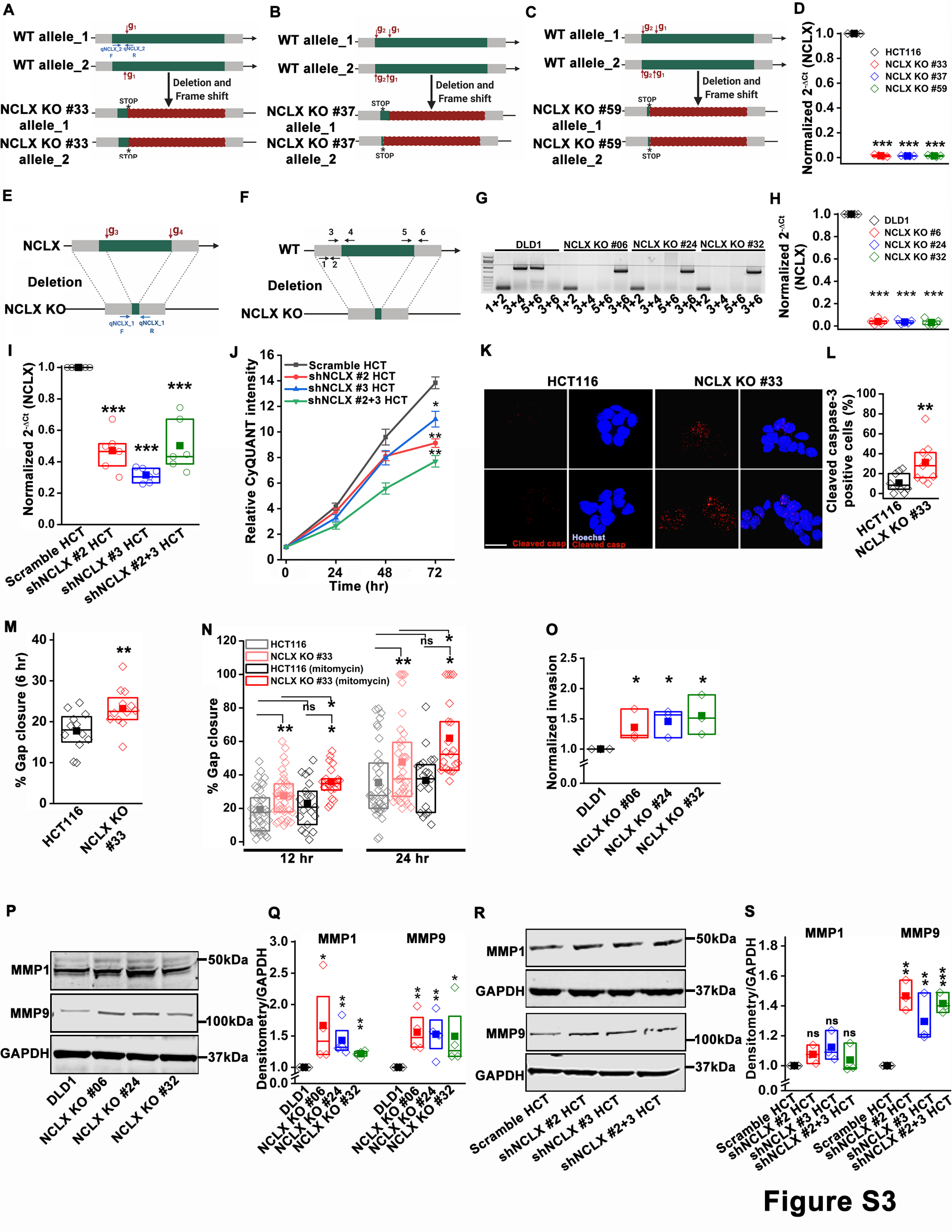

A
WT allele_1
WT allele_2
Deletion and Frame shift
NCLX KO #33 allele_1
NCLX KO #33 allele_2
STOP
B
NCLX KO #37 allele_1
NCLX KO #37 allele_2
C
NCLX KO #59 allele_1
NCLX KO #59 allele_2
D
Normalized 2-ΔCt (NCLX)
HCT116
NCLX KO #33
NCLX KO #37
NCLX KO #59
E
NCLX
Deletion
NCLX KO
F
WT
G
DLD1
NCLX KO #06
NCLX KO #24
NCLX KO #32
1+2 3+4 5+6 3+6
H
DLD1
NCLX KO #6
NCLX KO #24
NCLX KO #32
I
Scramble HCT
shNCLX #2 HCT
shNCLX #3 HCT
shNCLX #2+3 HCT
J
Relative CyQUANT intensity
Time (hr)
K
HCT116
NCLX KO #33
Cleaved casp
Hoechst
L
Cleaved caspase-3 positive cells (%)
M
% Gap closure (6 hr)
N
% Gap closure
HCT116 (mitomycin)
NCLX KO #33 (mitomycin)
12 hr
24 hr
O
Normalized invasion
P
MMP1
MMP9
GAPDH
50kDa
100kDa
37kDa
Q
Densitometry/GAPDH
R
S
Figure S3

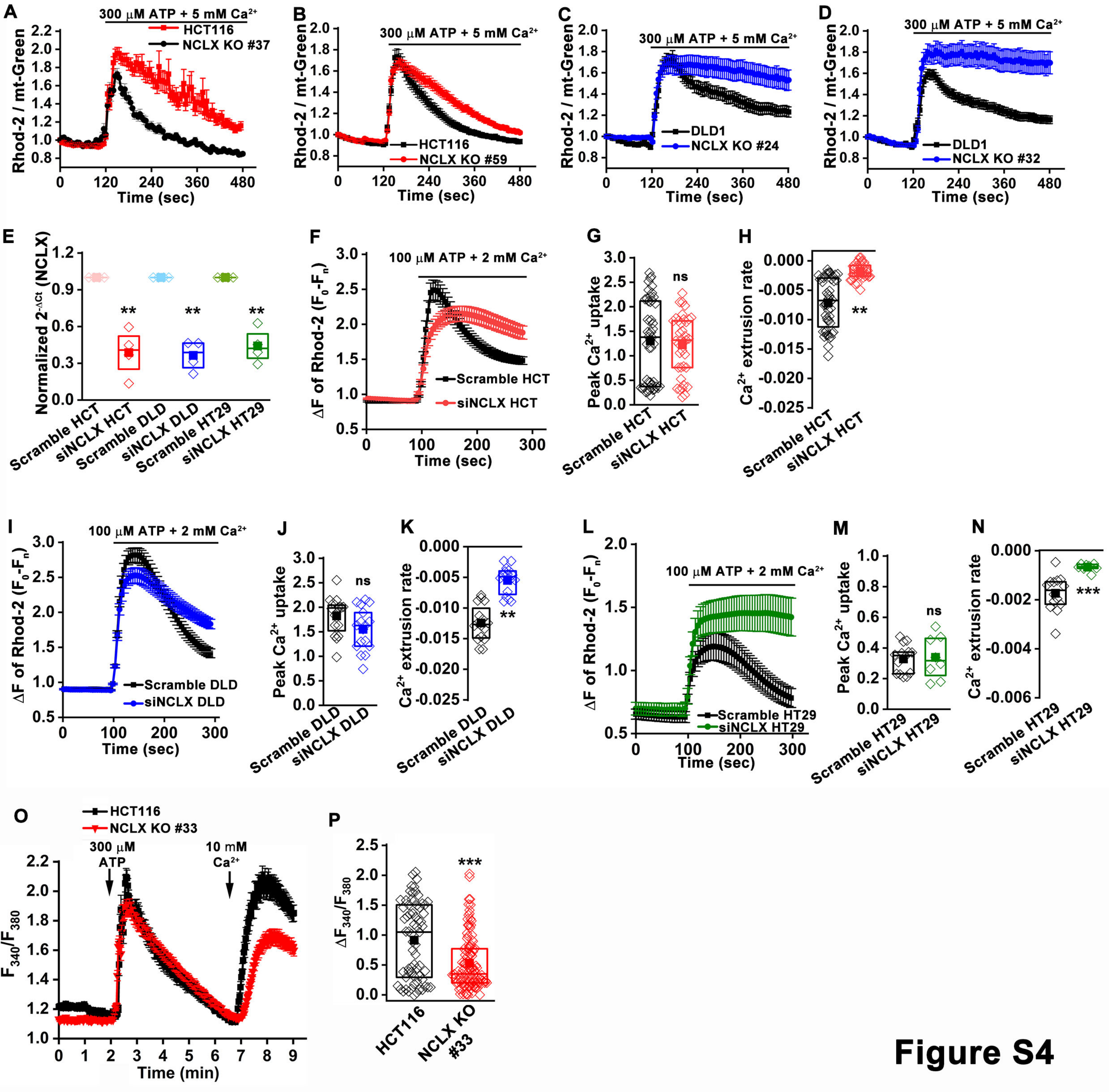

A
300 μM ATP + 5 mM Ca2+
HCT116
NCLX KO #37
Rhod-2 / mt-Green
Time (sec)
B
300 μM ATP + 5 mM Ca2+
HCT116
NCLX KO #59
C
300 μM ATP + 5 mM Ca2+
DLD1
NCLX KO #24
D
300 μM ATP + 5 mM Ca2+
DLD1
NCLX KO #32
E
Normalized 2-ΔCt (NCLX)
Scramble HCT
siNCLX HCT
Scramble DLD
siNCLX DLD
Scramble HT29
siNCLX HT29
F
100 μM ATP + 2 mM Ca2+
ΔF of Rhod-2 (F0-Fn)
Scramble HCT
siNCLX HCT
G
Peak Ca2+ uptake
ns
H
Ca2+ extrusion rate
I
100 μM ATP + 2 mM Ca2+
Scramble DLD
siNCLX DLD
J
K
L
100 μM ATP + 2 mM Ca2+
Scramble HT29
siNCLX HT29
M
N
O
HCT116
NCLX KO #33
300 μM ATP
10 mM Ca2+
F340/F380
Time (min)
P
ΔF340/F380
NCLX KO #33

### Figure S4

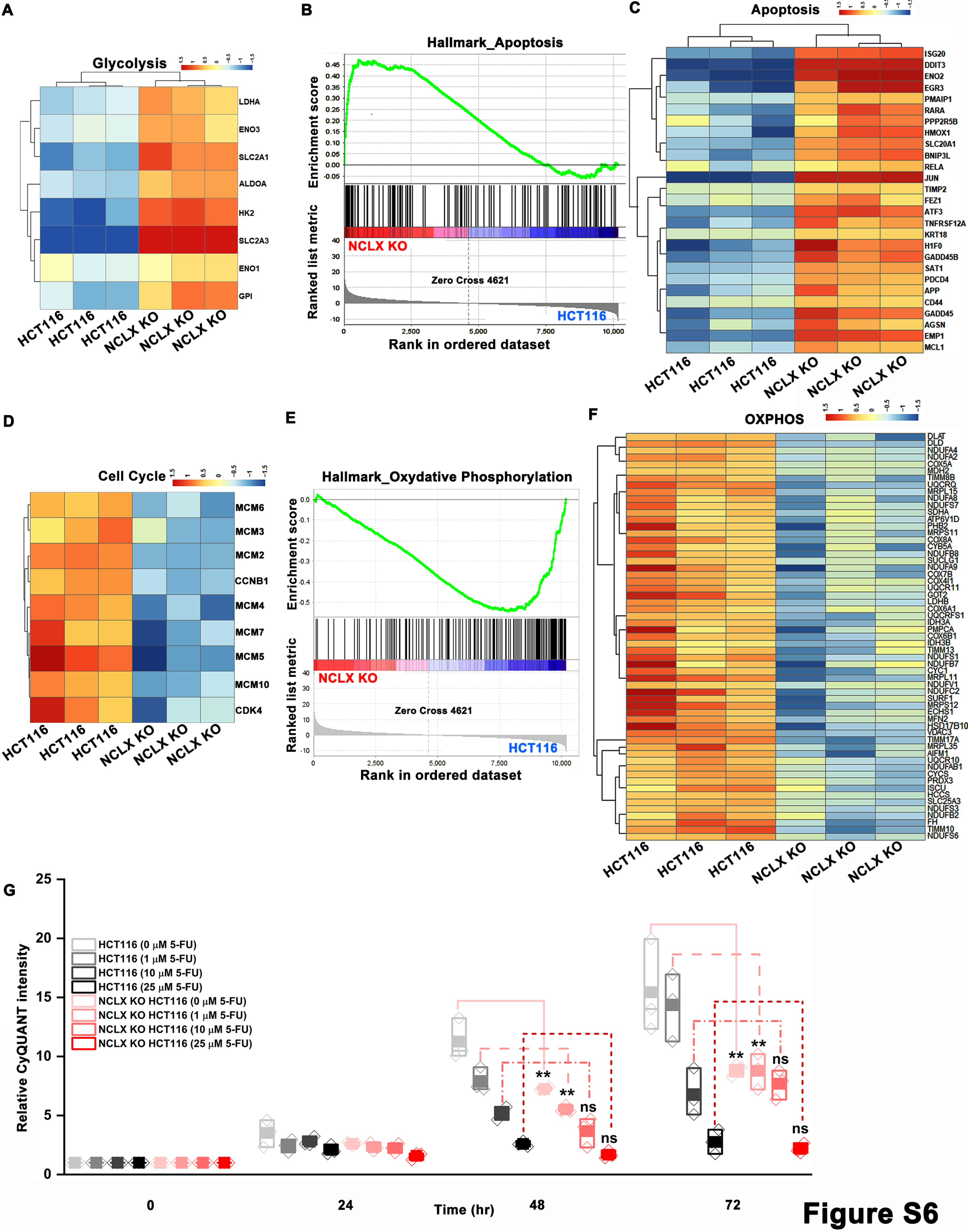
A
Glycolysis
HCT116
NCLX KO
LDHA
ENO3
SLC2A1
ALDOA
HK2
SLC2A3
ENO1
GPI
B
Hallmark_Apoptosis
Enrichment score
Ranked list metric
NCLX KO
Zero Cross 4621
HCT116
Rank in ordered dataset
C
Apoptosis
ISG20
DDIT3
ENO2
EGR3
PMAIP1
RARA
PPP2R5B
HMOX1
SLC20A1
BNIP3L
RELA
JUN
TIMP2
FEZ1
ATF3
TNFRSF12A
KRT18
H1F0
GADD45B
SAT1
PDCD4
APP
CD44
GADD45
AGSN
EMP1
MCL1
D
Cell Cycle
MCM6
MCM3
MCM2
CCNB1
MCM4
MCM7
MCM5
MCM10
CDK4
E
Hallmark_Oxydative Phosphorylation
F
OXPHOS
G
HCT116 (0 μM 5-FU)
HCT116 (1 μM 5-FU)
HCT116 (10 μM 5-FU)
HCT116 (25 μM 5-FU)
NCLX KO HCT116 (0 μM 5-FU)
NCLX KO HCT116 (1 μM 5-FU)
NCLX KO HCT116 (10 μM 5-FU)
NCLX KO HCT116 (25 μM 5-FU)
Relative CyQUANT intensity
Time (hr)
**
ns

Figure S6

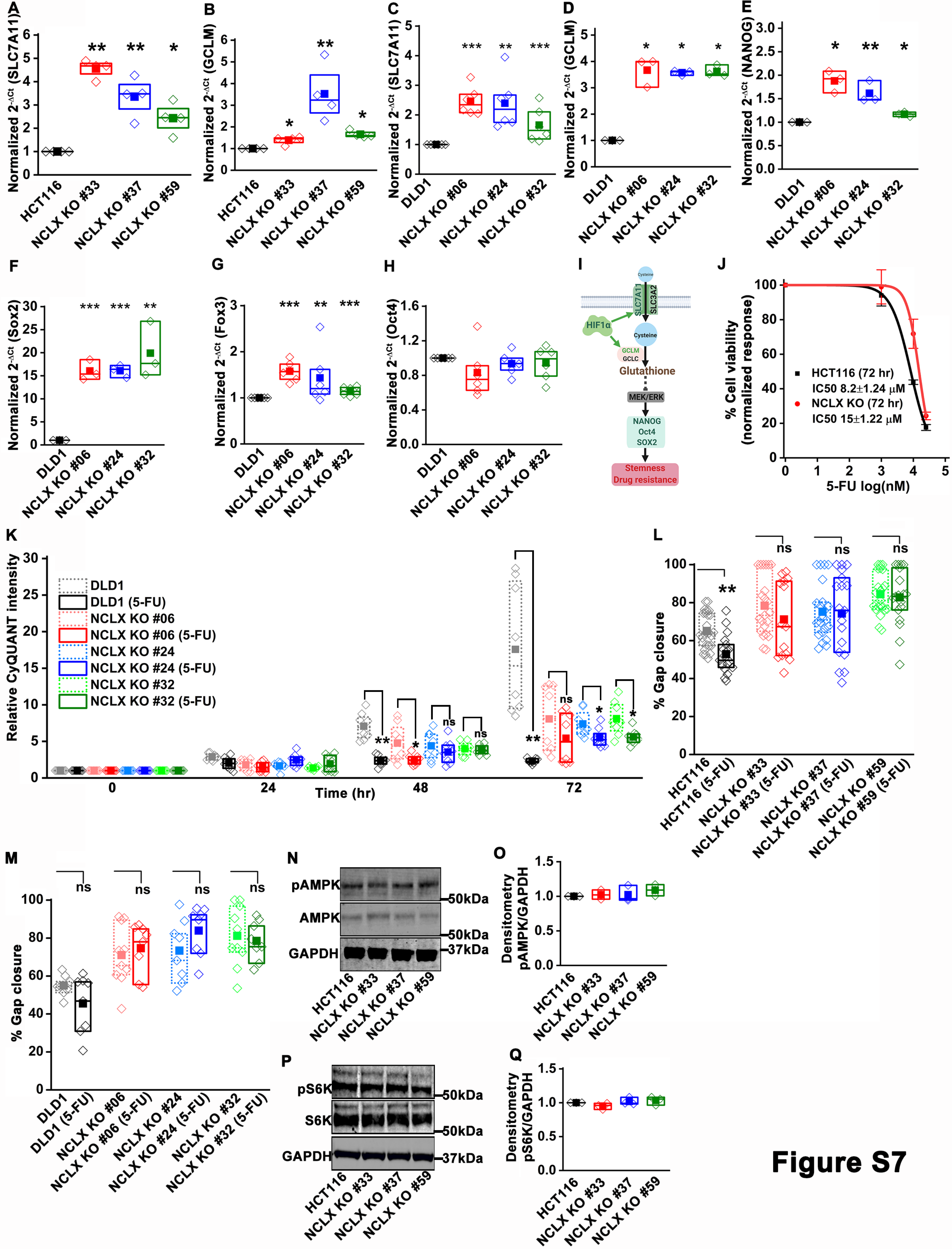
A
Normalized 2-ΔCt (SLC7A11)
HCT116
NCLX KO #33
NCLX KO #37
NCLX KO #59
B
Normalized 2-ΔCt (GCLM)
C
Normalized 2-ΔCt (SLC7A11)
DLD1
NCLX KO #06
NCLX KO #24
NCLX KO #32
D
Normalized 2-ΔCt (GCLM)
E
Normalized 2-ΔCt (NANOG)
F
Normalized 2-ΔCt (Sox2)
G
Normalized 2-ΔCt (Fox3)
H
Normalized 2-ΔCt (Oct4)
I
Cysteine
SLC7A11
SLC3A2
HIF1α
GCLM
GCLC
Glutathione
MEK/ERK
NANOG
Oct4
SOX2
Stemness
Drug resistance
J
% Cell viability (normalized response)
HCT116 (72 hr)
IC50 8.2±1.24 μM
NCLX KO (72 hr)
IC50 15±1.22 μM
5-FU log(nM)
K
Relative CyQUANT intensity
DLD1
DLD1 (5-FU)
NCLX KO #06
NCLX KO #06 (5-FU)
NCLX KO #24
NCLX KO #24 (5-FU)
NCLX KO #32
NCLX KO #32 (5-FU)
Time (hr)
L
% Gap closure
HCT116 (5-FU)
NCLX KO #33 (5-FU)
NCLX KO #37 (5-FU)
NCLX KO #59 (5-FU)
M
% Gap closure
NCLX KO #06 (5-FU)
NCLX KO #24 (5-FU)
NCLX KO #32 (5-FU)
N
pAMPK
AMPK
GAPDH
50kDa
50kDa
37kDa
O
Densitometry pAMPK/GAPDH
P
pS6K
S6K
GAPDH
Q
Densitometry pS6K/GAPDH

Figure S7

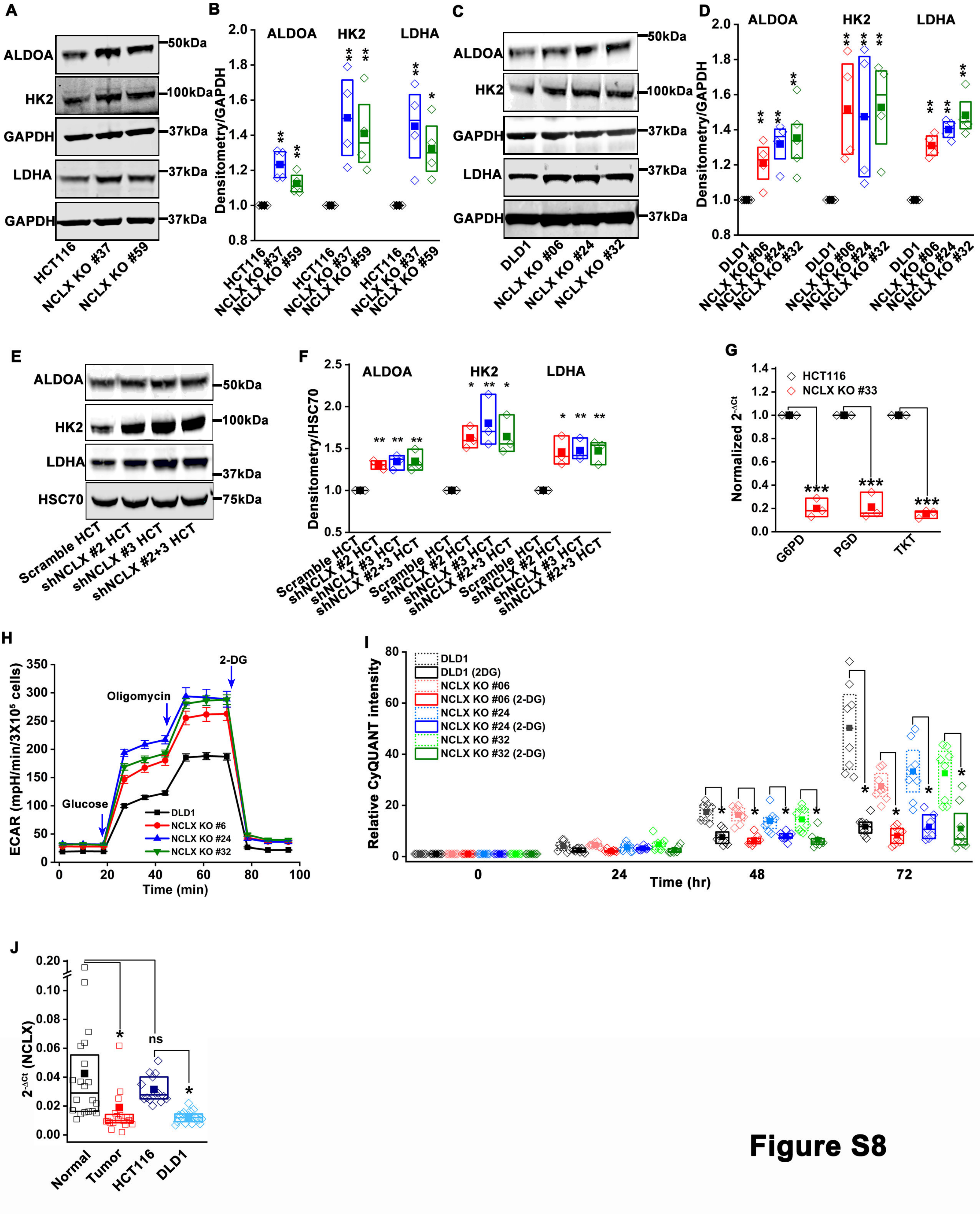
A
ALDOA
HK2
GAPDH
LDHA
GAPDH
50kDa
100kDa
37kDa
HCT116
NCLX KO #37
NCLX KO #59
B
ALDOA
HK2
LDHA
Densitometry/GAPDH
C
DLD1
NCLX KO #06
NCLX KO #24
NCLX KO #32
D
E
Scramble HCT
shNCLX #2 HCT
shNCLX #3 HCT
shNCLX #2+3 HCT
HSC70
75kDa
F
Densitometry/HSC70
G
HCT116
NCLX KO #33
Normalized 2^-ΔCt
G6PD
PGD
TKT
H
ECAR (mpH/min/3X10^5 cells)
Glucose
Oligomycin
2-DG
DLD1
NCLX KO #6
NCLX KO #24
NCLX KO #32
Time (min)
I
DLD1
DLD1 (2DG)
NCLX KO #06
NCLX KO #06 (2-DG)
NCLX KO #24
NCLX KO #24 (2-DG)
NCLX KO #32
NCLX KO #32 (2-DG)
Relative CyQUANT intensity
Time (hr)
J
2^-ΔCt (NCLX)
ns
Normal
Tumor
HCT116
DLD1

Figure S8
